# Improving Data Quality, Model Transparency and Performance in Lung Histopathology with Explainable AI

**DOI:** 10.64898/2026.09.14.751577

**Authors:** Salma Kazemi Rashed, Mikael Nilsson, Sonja Aits

## Abstract

Convolutional neural networks (CNNs) have shown strong capabilities for image analysis. However, deploying these models in medical settings is complicated by their limited transparency. Over recent years, many approaches have been developed to overcome the so-called “black box” problem of deep neural networks. Here, we show how such explainable AI (XAI) approaches can be applied to not only improve transparency but also training data quality and model performance, with classification of lung damage in histopathology images as use case. First, we conducted a thorough exploratory data analysis and visually compared the compressed multi-dimensional representations of the histology images from the last CNN layers with labels given by pathologists to reveal flaws in the training data. Second, we used Gradient-based Class Activation Mapping (Grad-CAM) as well as SHapley Additive exPlanations (SHAP) values to identify image regions that significantly contributed to the model decisions. To overcome identified shortcomings, we then finetuned additional top layers of the CNNs and introduced model architectures with attention which improved model performance. In summary, we developed a practical workflow that uses interpretability analyses to examine model perception, assess label consistency, and guide model refinement on the example of lung histopathology scoring, demonstrating how XAI approaches can increase both transparency and model performance. The code for this paper is shared at https://github.com/Aitslab/Histology_XAI.git.

## Introduction

In recent years, CNNs have gained prominence for medical and biological image analysis [1, 2]. They are already used in clinical trials or as tools to assist clinicians [3–5]. However, the “black box” nature of CNNs and other deep neural networks is a major impediment for building trust among healthcare professionals. It also complicates the approval of promising products for companies, with established regulation frameworks (e.g. from the Food and Drug Administration [FDA], and European Medicines Agency) still lacking entirely clear guidelines for usage of these models in formal clinical protocols [6, 7]. Although substantial progress has been made, widely accepted guidelines for the use of explainable medical AI have yet to be established, making it difficult to fully trust and standardize its application [8].

Digital histopathology is a dynamic field that has greatly benefited from automation using AI-based tools for histology applications [9, 10], However, these tools often struggle to provide truly trustworthy explanations for clinicians, even when efforts are made to improve their explainability [11–13]. Classification of scanned microscopic slides can be conducted faster and more accurately by CNN models compared to pathologists [14], but are only likely to attain widespread use if they are able to provide pathologists with the rationale behind the system’s decisions and conclusions in a simple way [15, 16]. With millions of parameters, CNNs can never be fully explainable, but several approaches have been developed to give insights toward the logic behind the decision made by the CNNs [17–19]. XAI for histopathological image analysis has employed several approaches such as IF-THEN rule-extraction, gradient- (e.g. Gradient-based Class Activation Mapping, Grad-CAM) and feature-based attribution methods (e.g. Shapley Additive exPlanations, SHAP), which can be used to visualize which image regions influence model decisions, or occlusion-based methods, which systematically mask image regions and observe changes in prediction scores [17, 18, 20–25]. Even though post-hoc explainability methods such as SHAP and Grad-CAM are widely used [26] they can be limited by their susceptibility to hyper-selectivity, whereby explanations emphasize only a subset of factors influencing a prediction. Other XAI approaches use attention-guided architectures to enhance both prediction and interpretability. Attention mechanisms can generate smooth, biologically coherent maps that align with expert annotations and highlight diagnostically relevant regions. While this can improve transparency by integrating interpretability into the learning process, attention-based approaches are often more computationally expensive and may require model-specific design innovations [27–29]. For all XAI methods, an important challenge is to connect the pixel patterns which represent known tissue morphologies and pathologies to the features extracted by CNN [30] because the model may rely on confounding features instead of true biomarker patterns or rely on true that are not readily interpretable by pathologists [31, 32]. XAI may thus misalign with clinical reasoning and even increase confusion and cognitive load among intended medical end-users, exposing a gap between technical transparency and practical utility [11, 33].

The need for explainability is especially desirable in difficult bioimage analysis tasks such as automated pulmonary histopathology scoring [34], where not only a variety of complex damage features needs to be considered (e.g. inflammatory cell influx into interstitial spaces, presence of proteinaceous debris within or thick membranes around alveolar spaces [35–37]) but also the subjectivity of pathologists in assessing these patterns [38]. Several studies have used CNNs to automate scoring of lung damage or lung cancer but used CNNs as “black-box”, without explaining the predictions made by the model [39–42]. More recent work has explored the application of XAI approaches for lung cancer histopathology, demonstrating their usefulness for understanding model behavior post hoc [43–51].

The present study takes this one step further by developing an iterative workflow which not only integrates multiple XAI approaches but also uses them to reveal data inconsistencies and guide model improvement through fine-tuning and architecture modification. We demonstrate the practical use of this workflow to improve existing models for lung damage scoring from our previous study [52], extending XAI approaches into a new histopathology domain.

Specifically, this work makes the following contributions:

1. We integrate complementary post-hoc XAI approaches, including latent-space visualization [53], Grad-CAM [54], and SHAP [26], into an XAI workflow and demonstrate its use on computer vision models for lung damage scoring
2. We show how these analyses can be used to assess the consistency and limitations of slide-derived tile labels and to relate learned model features to pathology-relevant tissue patterns.
3. We demonstrate how insights from these interpretability analyses can guide targeted model refinement through selective fine-tuning and attention-based architectures, leading to significantly improved state-of-the-art.
4. We incorporate an exploration of model complexity and inference time and relate it to performance into the workflow so it can also guide deployment decisions.

## Methods

### Use Case – Dataset and Original Models

We used our recent study about CNN-based histological scoring of lung damage [52] as an example use case to demonstrate the application of our iterative XAI workflow. In our previous work, we had trained CNN-based classification models to score the amount of lung damage in lung biopsy sections stained with Hematoxylin and Eosin (H&E) from a porcine model of acute respiratory distress syndrome (ARDS). The dataset is publicly available from https://www.ebi.ac.uk/biostudies/studies/S-BIAD419 [55, 56]. Sample groups included healthy control pigs or pigs subjected to lipopolysaccharide (LPS), an inflammatory agent used to model acute respiratory distress syndrome (ARDS), with or without mechanical ventilation (MV) or extracorporeal membrane oxygenation (ECMO) interventions, which are commonly used in ARDS patients.

The degree of lung damage had been scored by five annotators according to the Silva scoring system [57] which assesses seven damage features (Figure 1): A) Presence of inflammatory cells such as neutrophils, macrophages, and lymphocytes in interstitial spaces, B) presence of hyaline membranes (thick pink membranes lining the alveolar spaces) [35], C) proteinaceous debris in the alveolar spaces [37], D) thickening of the alveolar wall, E) enhanced injury (combining several features), F) haemorrhage and G) atelectasis, observed as obstruction of air spaces. Total damage scores, obtained by summing the individual feature scores, had been binned into “low”, “medium” or “high” damage labels for model training [52].

**Figure 1.**
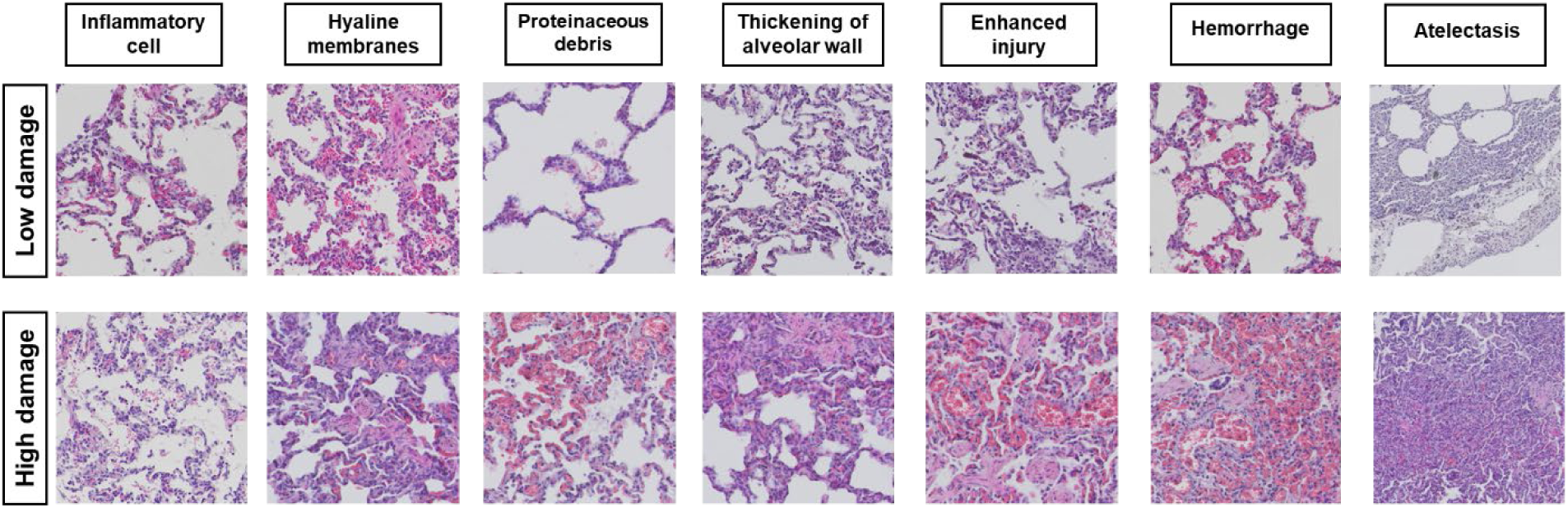
Damage features scored in the Silva scoring system. For each of the seven features examples of high and low damage are shown.

The dataset is relatively small, containing only 9–10 whole-slide images per treatment group, which is a common problem in medical research settings. No similar additional data could be obtained due to ethical, time and financial constraints. In addition, the images were very large, varied in dimension and contained sections with text and scale bars, which had to be removed. In the original study [52], we had addressed these issues by cutting images into small tiles (which increased sample number and removed text/bar-covered sections), with each tile inheriting the label of the parent image. In addition, 3-fold cross validation had been performed [52]. For this, tiles derived from the same animal were kept together in either training or evaluation sets, to avoid information leakage.

The two best individual models from the original study [52], which differed in architecture and augmentation strategy, were selected for further investigation in the present manuscript (Table 1). The V_r_a model is a VGG16 model [58] trained with augmentation consisting of mild rotation (–20 to +20 degrees), width and height shifting, flipping and change of brightness (0.8 to 1.3). The filling mode of empty border pixels formed by the shift or rotation had been set to ‘nearest’ pixels. The EN_m_a is an EfficientNetB4 model [59] with augmentation consisting of rotation (90, 180 and 270 degrees), flipping and change of brightness (0.8 to 1.3). Augmentation was performed using the Keras (version 2.8.0) “ImageDataGenerator” function with a custom rotation function for rotations 90, 180 and 270 degrees (script available at https://github.com/Aitslab/lunghisto).

**Table 1.**
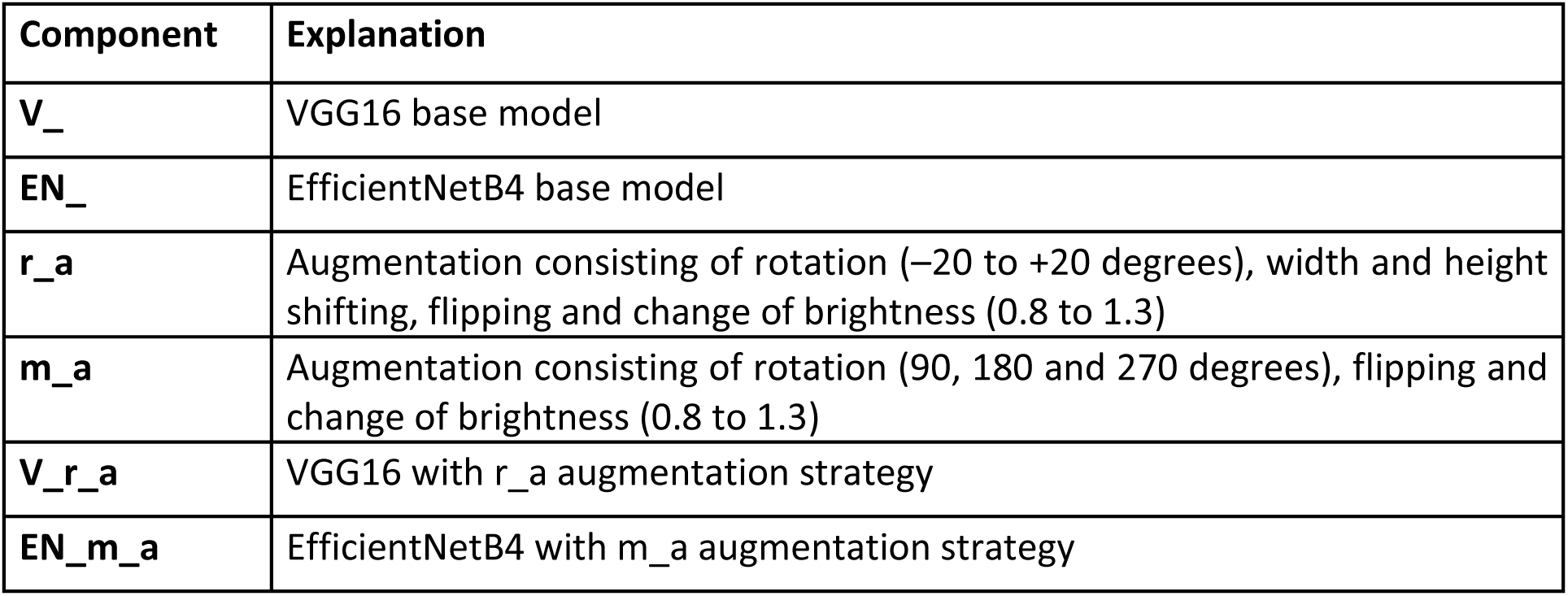
Model nomenclature. Derived from the original study [52].

| Component | Explanation |
| --- | --- |
| **V_** | VGG16 base model |
| **EN_** | EfficientNetB4 base model |
| **r_a** | Augmentation consisting of rotation (−20 to +20 degrees), width and height shifting, flipping and change of brightness (0.8 to 1.3) |
| **m_a** | Augmentation consisting of rotation (90, 180 and 270 degrees), flipping and change of brightness (0.8 to 1.3) |
| **V_r_a** | VGG16 with r_a augmentation strategy |
| **EN_m_a** | EfficientNetB4 with m_a augmentation strategy |

### Computational Procedures and Resources

New models (Table 2) were trained/fine-tuned on NVIDIA V100 and A100-SXM4-40GB GPU nodes of the Berzelius HPC cluster at the National Supercomputer Centre (NSC), Linköping University. In addition, we used a laptop running Windows with WSL and equipped with an NVIDIA RTX 2060 GPU (CUDA Version 11.8, 6.44 GB GPU Memory), 32 GB+ RAM System Memory, Keras 2.8.0 PyTorch 2.7.1 and Python 3.10.21 or higher for model training, XAI methods, inference time measurements, and other computations. The training scripts and hyperparameter settings used in the experiments are available in the accompanying code repository (https://github.com/Aitslab/Histology_XAI.git). Training procedures for all models were identical using 15-30 epochs, Adam optimizer and learning rate (lr=0.001-0.002).

**Table 2.** New model nomenclature, augmentations, and training datasets.

| Component | Explanation |
| --- | --- |
| **Data** | Same dataset as used in [52]. |
| **Data_7L** | A sevenfold enlarged variant of **Data** including blur augmentations (2×2 and 3×3 kernels), brightness shifts (0.8 and 1.3), and combined blur (using a 3x3 kernel)–brightness (0.8 and 1.3) transformations. |
| **V_c1** | VGG16 with Conv1 layer unfrozen – trained on **Data_7L**, along with “**m_a**” augmentation |
| **V_c1_c2** | VGG16 with Conv1 and Conv2 layers unfrozen – trained on **Data_7L**, along with “**m_a**” augmentation |
| **EN_BN** | EfficientNetB4 with Batch Normalization layer unfrozen – trained on **Data_7L**, along with “**m_a**” augmentation |
| **EN_BN_c1** | EfficientNetB4 with Batch Normalization and Conv1 layers unfrozen – trained on **Data_7L**, along with “**m_a**” augmentation |
| **V_m_a_Attn** | VGG16 + Attention, trained on **Data**, along with “**m_a**” augmentation |
| **EN_m_a_Attn** | EfficientNetB4 + Attention, trained on **Data**, along with “**m_a**” augmentation |
| **ViT (tiny)** | vit_tiny_patch16_224 (192 hidden layers), trained on **Data** normalized to ImageNet mean and standard deviation |
| **ViT (small)** | Vit_small_patch16_224 (384 hidden layers), trained on **Data** normalized to ImageNet mean and standard deviation |
| **V_Attn** | VGG16 + Attention, trained on **Data** normalized to ImageNet mean and standard deviation |
| **EN_Attn** | EfficientNetB4+ Attention, trained on **Data** normalized to ImageNet mean and standard deviation |

### Unsupervised Clustering of Latent Representations

For unsupervised clustering, the tiles from the evaluation sets in each fold were passed to the V_r_a and EN_m_a models for prediction. The feature vectors generated for each prediction, which had a length of 128, were extracted from the final fully connected dense layer (Figure 2). Three techniques for dimensionality reduction, Principal Component Analysis (PCA) [60], T-Distributed Stochastic Neighbor Embedding (t-SNE) [61], and Uniform Manifold Approximation and Projection (UMAP) [62], were then applied to feature vectors and original tiles (converted into vectors), followed by visualization in a 2D plot together with the predicted labels, the ground-truth labels, and the original histology slides. For PCA and t-SNE we used the scikit-learn library [63], and for UMAP the umap library [64]. With PCA (n=5), t-SNE and UMAP were run with their default settings (https://github.com/Aitslab/Histology_XAI/tree/main/Unsupervised).

**Figure 2.**
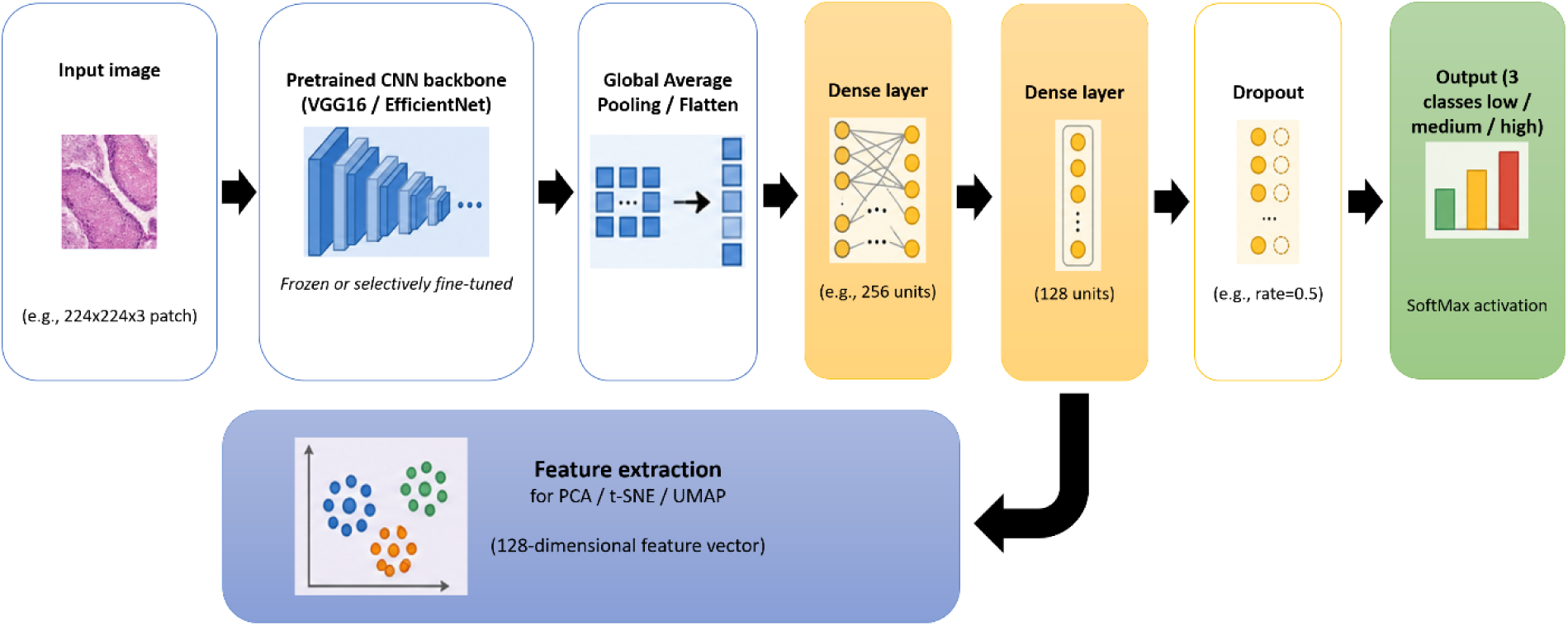
Extraction of latent representations for unsupervised clustering. A compressed representation of the input was obtained from the final fully connected dense layer. For each prediction, a 128-dimensional feature vector was extracted.

### Grad-CAM

The Grad-CAM [54] algorithm calculates the gradients of each target class (e.g., ‘low’, ‘medium’, and ’high’) with respect to the last convolutional layer of the model, highlighting areas of the image which influence the final prediction of that class. The highlighted map is a coarse-grained activation map which needs to be resized to be able to show the activated areas of the original input image. The pseudo code behind the Grad-CAM heatmaps is summarized in Algorithm I [54]. Briefly, to generate a Grad-CAM heat-map, we start by creating a model with probes added to the layers where we want to generate the heat-map. We then attach fully connected layers to the core model for prediction and pass the input image through the model to capture the feature maps of the target convolutional layer and the logits before the SoftMax layer. Next, we compute the gradient of the the predicted-class logit with respect to the feature maps of the target layer. We then calculate the importance weight for each feature map channel by globally averaging the gradients over the spatial dimensions. These weights are used to compute a weighted sum of the feature maps. We take the positive part of the result, resize it to match the input image size, and use it to generate the heat-map (script at https://github.com/Aitslab/Histology_XAI/main/GradCAM).

#### Algorithm I

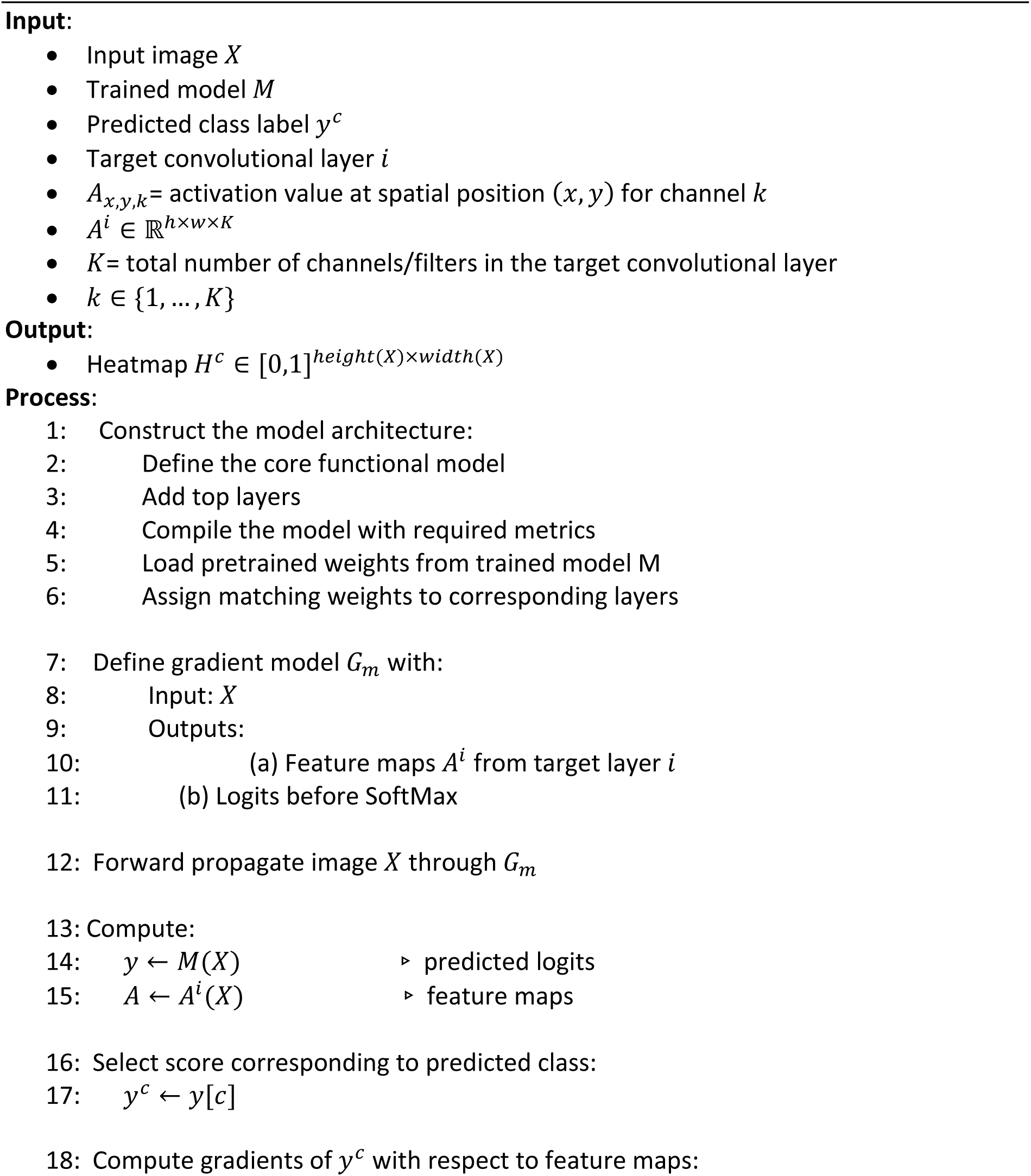
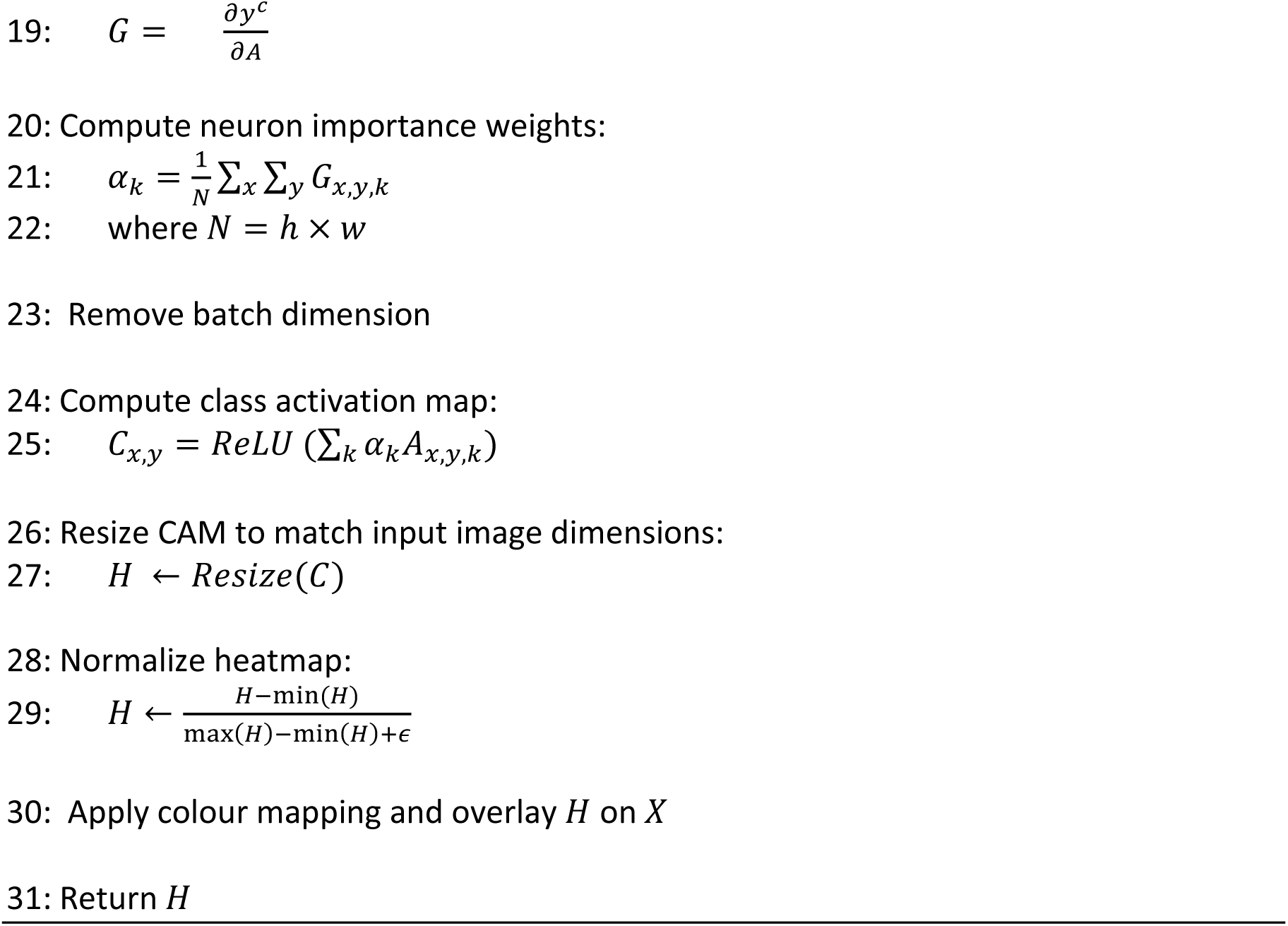

### Replacement of Activation Function

While computing Grad-CAMs, we observed issues in some tile maps, which we traced back to the ReLU activation function causing a large portion of features to be zeroed out. We therefore replaced the ’ReLU’ function with “ELU”, “LeakyReLU”, “LeakyReLU-ReLU” functions, which allow maintenance of gradient flow and ensure that more tiles contribute to the final prediction, and then re-trained the model and evaluated the new models on the validation sets.

### SHAP

SHAP values [26] are used to assess the significance of individual features towards a particular output, such as output of class *c* in a classification model, where the general idea was to consider a model as a game where data points (pixel values in image data) act like players in cooperation with other features to make the output [26, 65]. The SHAP value is computed by considering all possible coalitions of features that contribute to the classification decision and measuring the contribution of each pixel/feature to each set of features. Computing the SHAP values for every pixel/feature can be resource intensive. Thus, approximate methods can provide a reasonable estimate for a subset of features (regions) and help identify the most influential regions of the data for a given decision. We used the explainer function (shap.Explainer), which combines the model with a masker that masks the withheld features for each coalition of features, resulting in a matrix of SHAP values associated with their respective credits (positive or negative) in the final assignment of that input to each class. The pseudo-code for computing the SHAP value for is shown as Algorithm II (script at https://github.com/Aitslab/Histology_XAI/main/SHAP).

#### Algorithm II

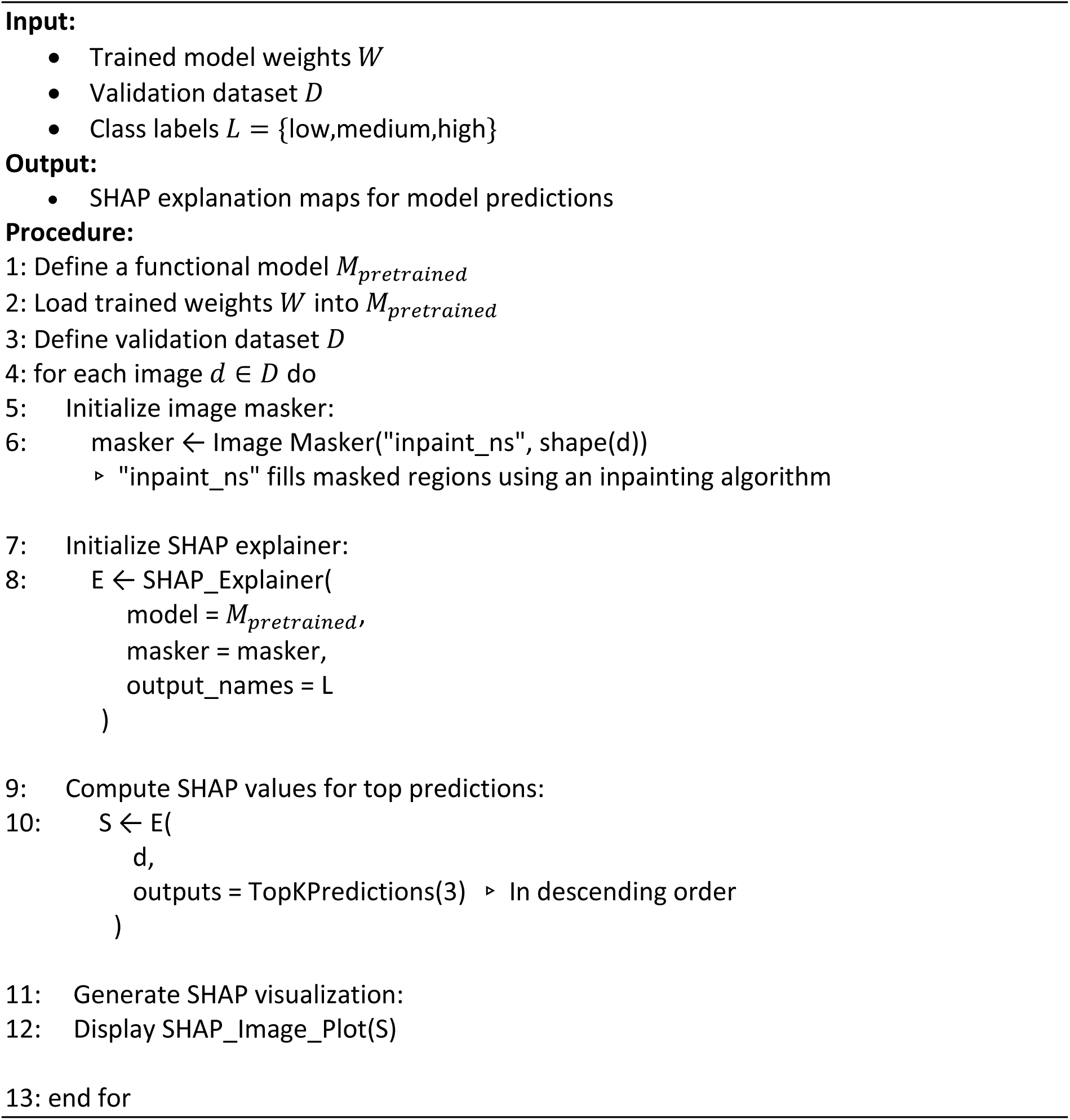

### Deeper Model Finetuning

To improve interpretability as well as assessing whether explainability-driven adjustments could lead to better model performance, we fine-tuned additional layers (Figure 3), thereby altering both the number and type of trainable parameters. We unfroze one or two convolutional layers for the VGG16 model (V_c1, V_c1_c2) and batch normalization and convolutional layers for EffNetB4 model (EN_BN, EN_BN_c1), respectively (Figure 3) by setting those layers as trainable in the training script.

**Figure 3.**
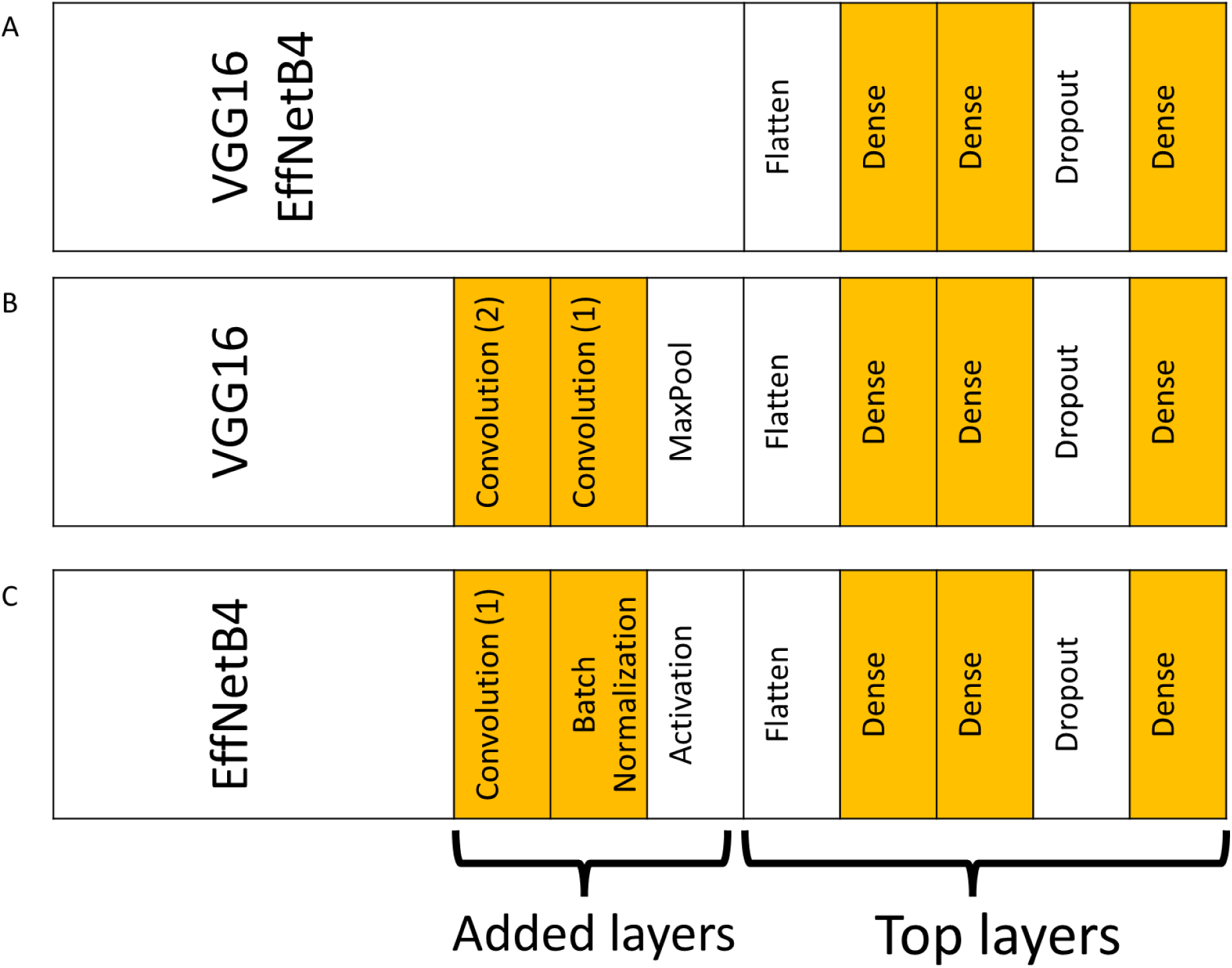
Overview over original and newly trained models. A) The original “V_r_a” and “EN_m_a” models with the core model and trainable top layers, B) The V_c1 and V_c1_c2 models with newly trainable convolutional layers (1) and (2). C) The EN_BN, and EN_BN_c1 models with newly trainable batch normalization and convolutional layers (1). Yellow cells indicate trainable layers, while white cells represent layers without trainable parameters.

To avoid overfitting when increasing the number of trainable parameters, we expanded the training dataset sevenfold by adding deterministic augmentations, namely brightened and blurred versions of the images (blurring filters with [2,2] and [3,3] kernel sizes, adjusting brightness to 0.8 and 1.3, and combinations of both blurring with kernel size of [3,3] and change of brightness to 0.8 and 1.3) (Figure 4 and Table 2) by ensuring that different versions of each image (e.g., blurred, brightened, and blurred + brightened) were included in the data pool. In addition, during training, all images undergo random augmentations through the “ImageDataGenerator” pipeline (Table 2), which further increases data variability and improves model robustness (https://github.com/Aitslab/Histology_XAI/tree/main/Train_more_layers/training_scripts).

**Figure 4.**
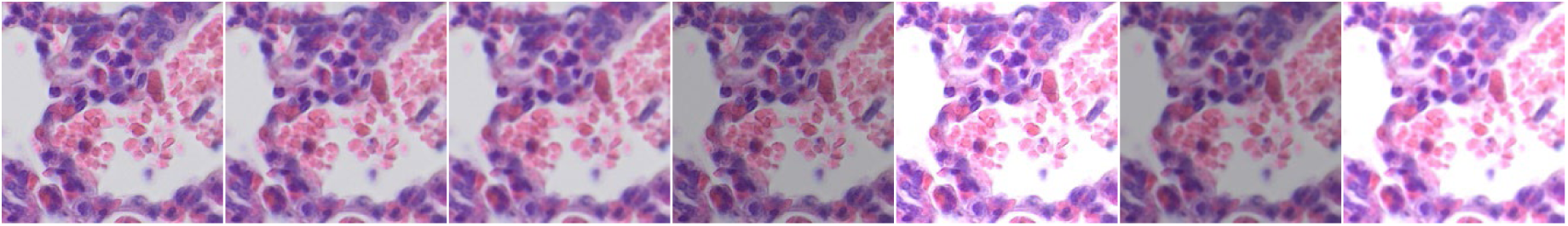
Example of blurring and brightness augmentation. Original image tile to the left. The augmentations applied sequentially from left to right include blurring with kernel sizes of (2,2) and (3,3), adjusting brightness to 0.8 and 1.3, and combinations of both blurring and brightness adjustments using a (3,3) blurring kernel with brightness ratios of 0.8 and 1.3.

Layer unfreezing during the training phase, along with the alignment of the saved model to the original network architecture, is performed according to Algorithm III.

#### Algorithm III

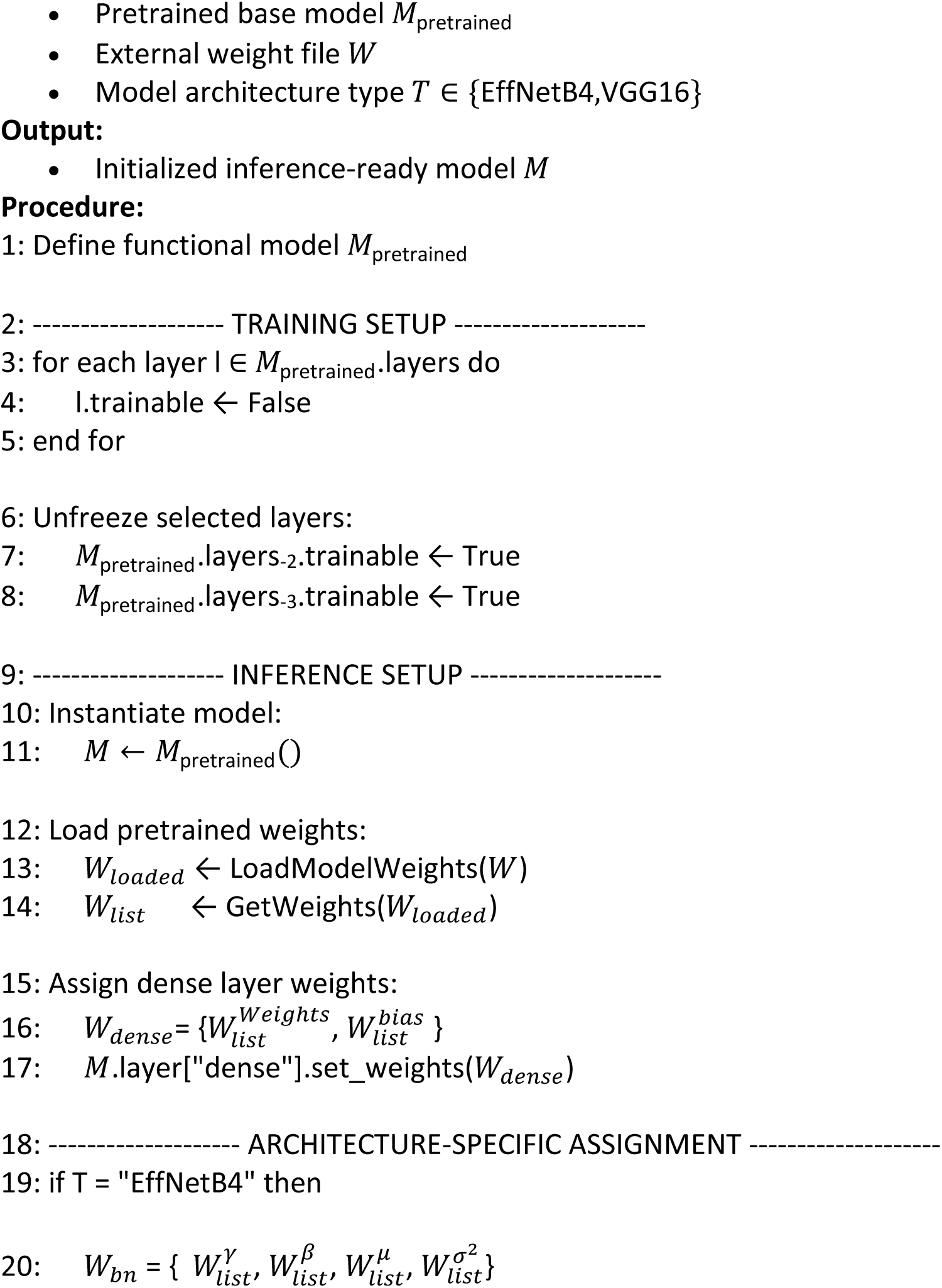
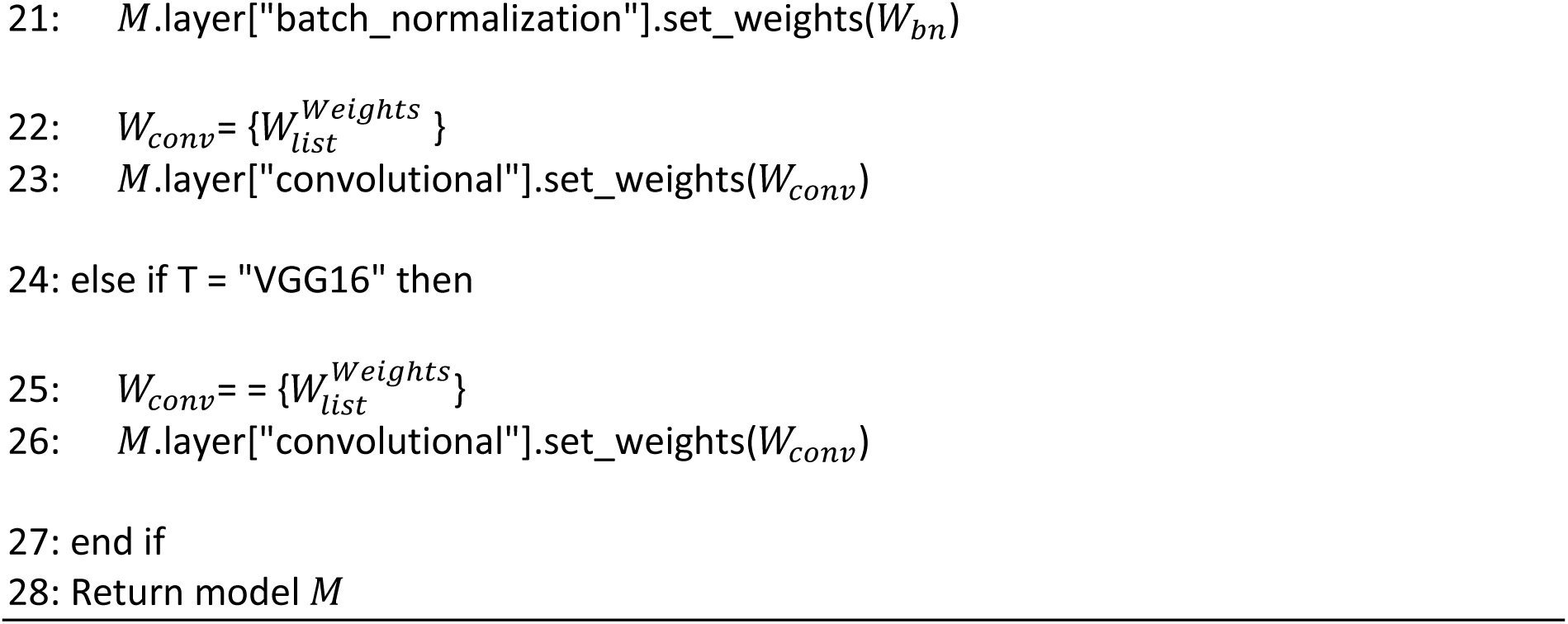

### Addition of Convolutional Attention Layers

To track performance with attention-driven features during training, we added convolutional and locally connected layers with an attention mechanism to the models (V_m_a_Attn, EN_m_a_Attn (Figure 5)). The convolutional layers were utilized as filters to extract features from images. This then converted to attention weights by a ‘Locally Connected’ layer with ‘sigmoid’ activation function. The attention weights then scaled and multiplied with image data. The masked features and the scaled attention weights were tracked in each epoch of training. Selected attention maps for the best model for each fold were inspected manually (i.e., selectively chosen). The attention models (V_m_a_Attn, EN_m_a_Attn) were trained on the same dataset as V_r_a and EN_m_a using the m_a augmentation strategy (Table 1). The same training script, number of epochs, and evaluation metrics were used across all experiments and are provided in the accompanying code repository (https://github.com/Aitslab/Histology_XAI/blob/main/Attention/Training).

**Figure 5.**
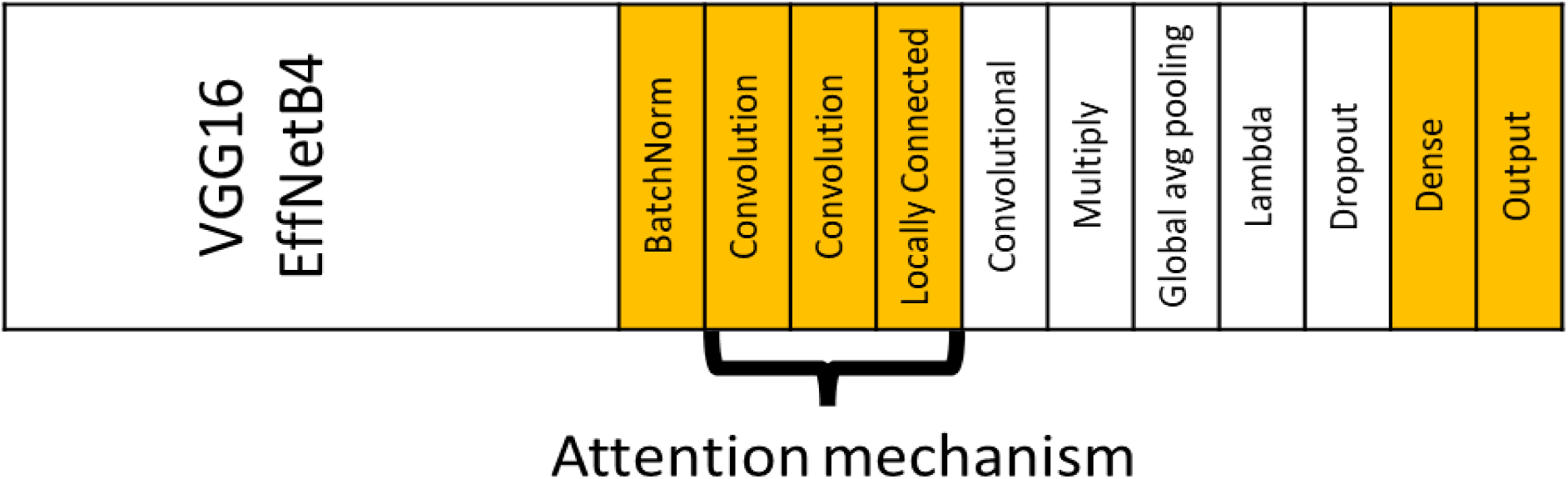
Addition of attention to the model architecture. This diagram depicts the V_m_a_Attn and EN_m_a_Attn models with attention mechanisms added to the pretrained based models. As before, orange layers are trainable layers where white layers are non-trainable layers (Frozen or no parameters). Layers are 1. VGG16 or EfficientNetB4 core (Trainable params: 0), 2. BatchNormalization (Trainable), 3. Attention Mechanism (3.1. First attention convolutional layer (Trainable), 3.2. Second attention convolutional layer (Trainable), 3.3. LocallyConnected2D (Trainable), 4. Up-projection convolutional layer (Trainable: 0 (explicitly frozen), 5. Multiply (no parameters), 6. GlobalAveragePooling2D (no parameters), 7. Lambda layer (no parameters), 8. Dropout (no parameters), 9. Dense (Trainable), 10. Output Dense (Trainable).

### Training of ViT Models

ViT models were adapted from a study by Amato et al. (https://github.com/salvatorecalderaro/Explainable-Histopathology-Classification-ViT) [51]. Briefly, we converted the architecture from the original binary classification to 3-class classification. Two ViT variants (tiny and small) were then trained on the lung damage dataset (https://github.com/Aitslab/Histology_XAI/tree/main/ViT) in a similar manner as the CNN-based models but using the original training scripts. This script also governed fold composition, which differed from the CNN models by randomly distributing tiles from each group across folds and not keeping images from the same pig together. Tiles were normalized using the ImageNet mean and standard deviation. No augmentation was applied. For direct comparison, we also trained two new CNN models (V_Attn and EN_Attn) with the same architecture as V_m_a_Attn and EN_m_a_Attn but using the fold composition, data preprocessing and training procedures of the ViT models.

### Statistical Analysis and Multi-dimensional Comparison

To determine whether the observed performance changes were statistically significant, we performed McNemar’s test, which relies on the comparison of paired predictions. We compared predictions on all validation samples across the three folds (8,073 total) by the modified V-type models (V_r_a (‘LeakyReLU_ReLU’, V_c1, V_c1_c2, V_m_a_Attn) versus the V_r_a baseline model and by the modified EN-type models (EN_BN, EN_BN_c1, and EN_m_a_Attn) versus the EN_m_a baseline model.

To judge the contribution of individual modifications we compared the different model configurations with regards to accuracy, F1-score, trainable parameter count, and inference time.

Accuracy and F1 scores were calculated on the respective evaluation set of each fold and then averaged across folds, where

Accuracy = (TP + TN) / (TP + TN + FP + FN),

F1 = 2 × (Precision × Recall) / (Precision + Recall),

Precision = TP / (TP + FP),

Recall = TP / (TP + FN),

where TP, TN, FP, and FN denote true positives, true negatives, false positives, and false negatives, respectively.

Inference times were measured on a laptop with a NVIDIA RTX 2060 GPU (see additional specifications under “Computational Procedures and Resources”).

## Results

### Lung damage histology scoring as use case for XAI workflows

Histology scoring is a complex medical image analysis task which is widely used in both research and clinical settings, making it an important domain for XAI. We chose our recent study about CNN-based histological scoring of lung damage [52] as an example use case for developing a workflow for XAI methodology. In this study, CNNs had been trained to categorize the overall lung damage on histology images from a porcine model of ARDS with different treatments into three levels of damage (low, medium, and high). The two best models from the study, called V_r_a and EN_m_a, relying on a VGG16 and EfficientNetB4 backbone, respectively, were chosen to be investigated in more detail to understand the model decisions. We also wanted to explore two issues with the ground truth-labels of this study, which may have confused the models during training and affected model evaluation. First, while pathologists were only able to perform scoring on whole-slide level, models were trained to assign labels to smaller image tiles, with each tile inheriting the label of the parent. This enabled a more detailed view of the lung damage but introduced potential label errors due to the spatial heterogeneity within slides. Second, problems with inter-annotator and intra-annotator reproducibility (not clearly linked to annotator experience) had been observed when examining correlations between individual scores and repeat scoring rounds undertaken by the same annotator [52]. This was not unexpected due to the difficulty of manual histology scoring but made the ground truth somewhat unreliable even though it was based on the mean and median scores of five annotators [52].

We designed an XAI workflow aimed at understanding and reducing data and model issues through several complementary steps: 1. Unsupervised Clustering of Latent Representations, 2. Grad-CAM, 3. SHAP values, 4. Deeper layer training, and 5. Introduction of attention-based architectures.

### Discovering Structure in Latent Representations via Unsupervised Clustering

To explore the performance of the models irrespective of potentially wrongful labels, we examined the predictions by V_r_a and EN_m_a models using unsupervised approaches. For this, we extracted the latent features representing the images from the final layers of deep convolutional neural networks Using PCA (n = 5), with explained variance ratios of [0.3771, 0.2477, 0.1245, 0.0708, and 0.0301], respectively, approximately 85% of the total variance was retained. We then reduced the features to two dimensions using PCA, t-SNE, and UMAP for visualization, displaying them alongside the predicted labels produced by the models, the ground-truth labels obtained from pathologists, and the original histology slides. This allowed us to explore how the latent space features, particularly from the last CNN layers, capture underlying patterns, and how these patterns appear when projected into a lower-dimensional space [53].

For the original tiles, no separate clusters were observed when applying PCA, t-SNE or UMAP (data not shown). When clustering the feature vectors (extracted from last dense layer) from each model (Figure 6), we observed distinct clusters for each fold. For the “V_r_a” model the fold 2 cluster differed in appearance, being smaller with PCA and split into subclusters using t-SNE and UMAP. We therefore examined fold differences in the feature vectors from this model in more detail. When visualizing the feature vector Euclidean norms (Supplemental Figure 1), we observed a shift in the values in fold 2 compared to the other two folds, with many feature vectors having a value of 0, causing some data points to be zeroed out by the activation function. This indicated differences in data distributions across folds.

**Figure 6.**
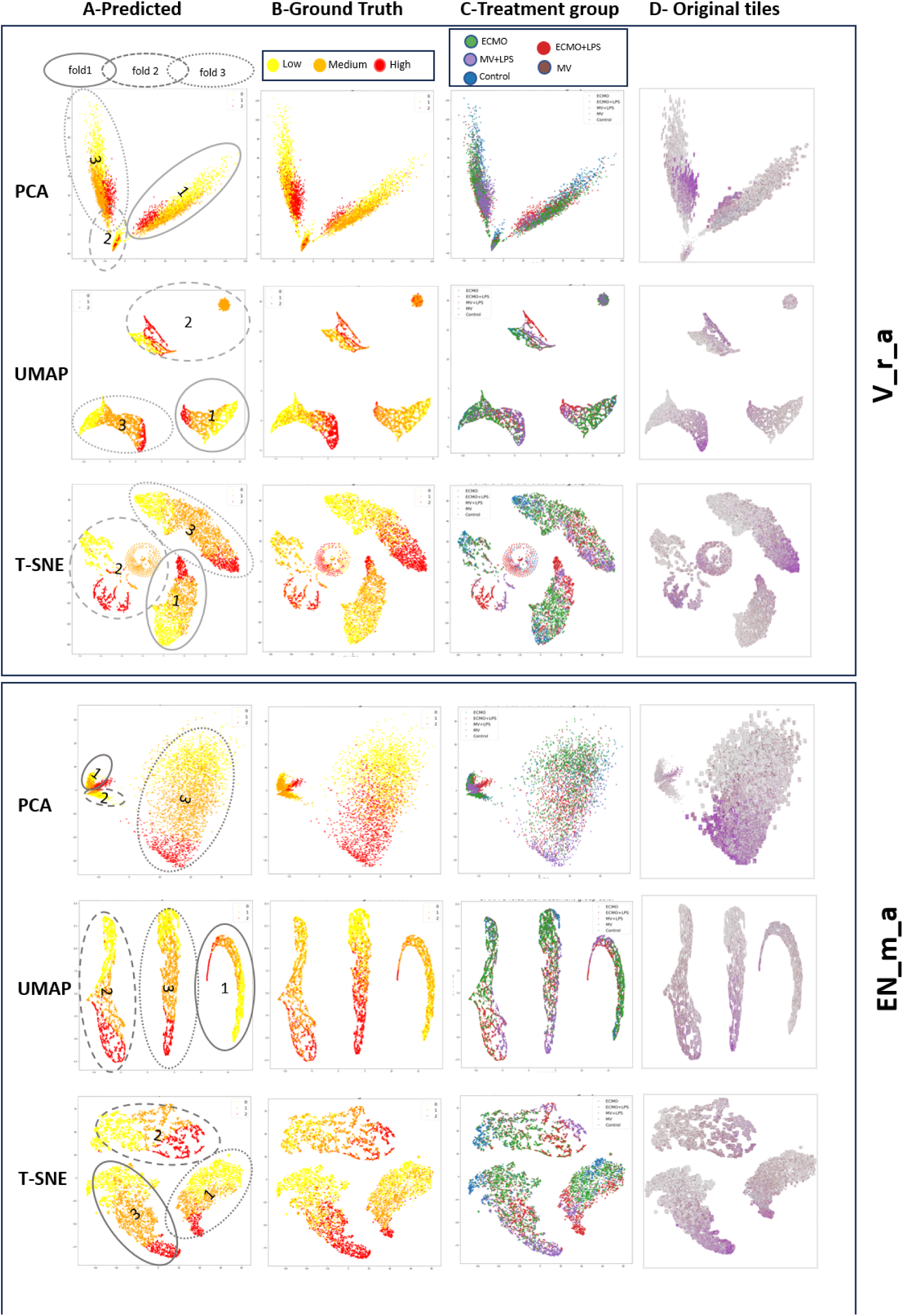
Unsupervised clustering of latent space vectors. Representation of latent space vectors of all datapoints (tiles) in 2D for all folds (where Folds are indicated by ellipses) and with colours labelled with (A) predicted scores by V_r_a and EN_m_a models, B) ground truth scores, (C) treatment groups and (D) original miniature image, where tiles with dark purple colour mostly contain areas of high damage.

Next, we compared the distribution of the predicted versus the ground-truth labels for both models by visualizing the labels on the clustering plots with distinct colors (Figure 6A, Figure 6B). In general, low, medium, and high labels were grouped within the clusters for both predicted and ground-truth labels, with low and high labels separated from each other by the medium labels. This separation was clearer in the predicted labels. The fully separated subclusters in fold 2 for the “V_r_a” model also corresponded to distinct predicted labels, whereas they had mixed ground-truth labels.

There are two explanations for the observed differences between predictions and ground truth. Either the models could have made errors, or the annotations were incorrect due to spatial heterogeneity or human annotation errors. To further explore which of these possible explanations might be correct, we exchanged the label-based colouring of the clusters with colouring corresponding to the treatment groups (Figure 6C) and original images of each tile (Figure 6D). When overlaying the miniature original tiles, we observed a distinct shift in colour from dark to very light from one side of the clusters to the other. Light colour typically corresponds to tiles with healthy tissue, which has wider airspaces (which appear white) and thin alveolar walls. In contrast darker purple/pink colouring reflects various types of damage, for example influx of inflammatory cells (dark purple), hyaline membrane-lined alveolar walls (pink), or atelectasis (collapse of airspaces reflected in loss of white areas in the image). The light–dark purple distribution is consistent with both the ground truth and predicted values in the high and low damage classes. However, in the medium damage class, the distribution shows a stronger resemblance to the predicted scores.

The treatment group labelling suggested that the damage was unevenly distributed across the slides with undamaged and damaged tiles generated from the same sample. When comparing the labelling of the treatment group to the ground truth labels, high-damage groups like LPS-induced samples treated with MV (MV+LPS) and ECMO (ECMO+LPS) almost perfectly matched with high damage datapoints in the ground truth plot. Other treatment groups such as ECMO, MV, and even control were scattered across yellow (low damage), orange (medium damage), and even red (high damage) data points. This supported the notion that tiles from the same treatment group displayed different levels of damage despite having the same ground truth score. This discrepancy may arise from the possibility that the treatments themselves cause lung injury, which was one of the central questions being assessed in the original ARDS study.

### Grad-CAM

Next, we used gradient-based algorithms, a widely used XAI method which tracks gradients of loss function values of the predicted class with respect to the last or deeper convolutional layers. It allows the visualization of activated regions of the image in that layer, which led to the prediction of the respective class, in a coarse heatmap [54].

After fine-tuning, we generated Grad-CAMs for the “V_r_a” and “EN_m_a” models (Figure 7, top block). These showed that, for certain predictions, especially in fold 2, some tiles were not activated for the predicted class and therefore did not contribute to the prediction. These tiles mostly had feature vector with a value of 0. The ReLU function, which was used as the activation function in the trainable layers of both models, outputs zero for negative input values. This can lead to the well-known "dying ReLU" problem, where neurons become inactive and cease contributing meaningful information to downstream layers. In contrast, other activation functions like LeakyReLU and Exponential Linear Unit (ELU) allow maintenance of gradient flow and ensure that more tiles contribute to the final prediction. We therefore replaced the ’ReLU’ function with “ELU”, “LeakyReLU”, “LeakyReLU-ReLU” functions, re-trained and evaluated the new models on the validation sets (Supplemental Figure 1, Figure 7, bottom block). Models which used ReLU-based activation functions (plain ReLU, or “LeakyReLU–ReLU”) for the last two fully connected layers, showed issues with disregarded tiles. Models with other activation functions showed improved activation patterns.

**Figure 7.**
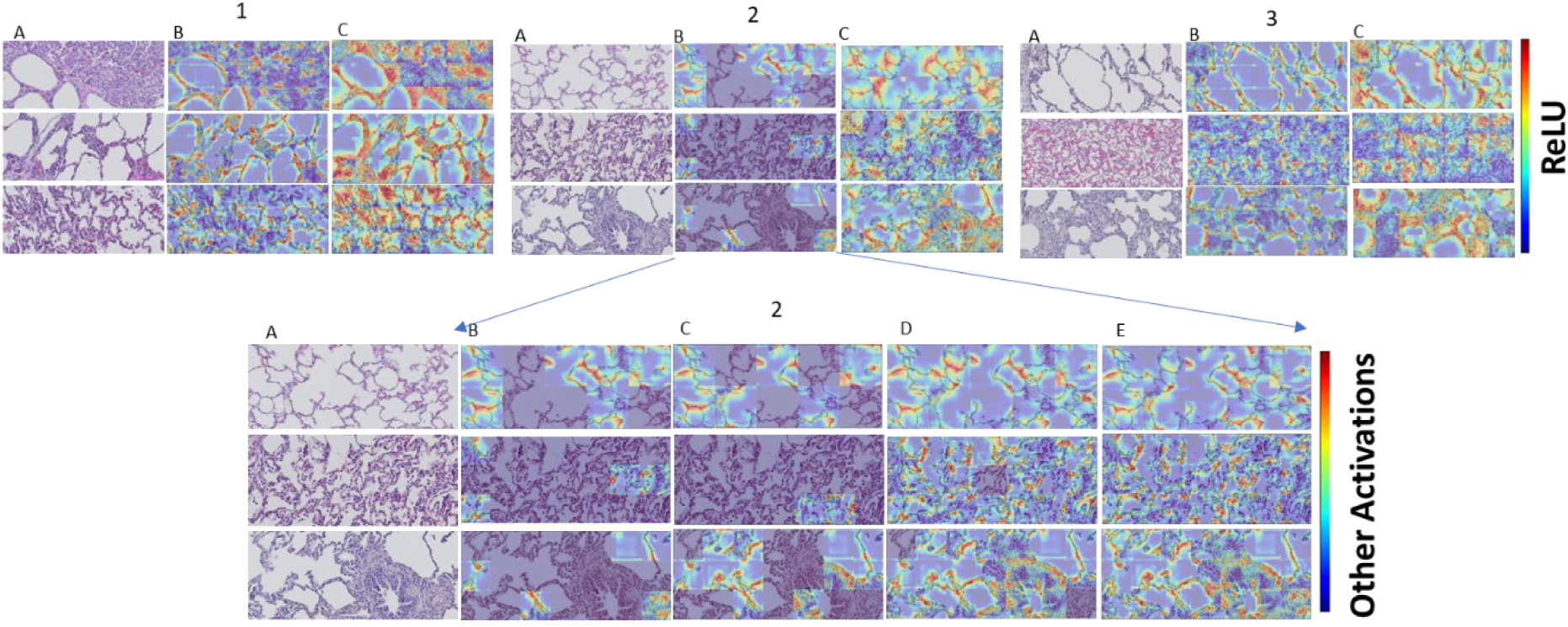
Class Activation Maps and Effect of Different Activation Functions. Top block - Samples of original images from folds 1, 2, and 3 (column A), the activation heatmaps shown on the original images coming from the “V_r_a” model (column B), and the activation heatmaps shown on the original images coming from the “EN_m_a” model (column C). Bottom Block-Samples of images from fold 2 (column A), the activation heatmap shown on the original images coming from the “V_r_a” model with B) ‘ReLU’, C) ‘LeakyReLU (0.2)_ReLU’, D) ‘ELU’ and E) ‘LeakyReLU’ activation functions. Non-activated tiles appear uniformly dark.

The altered activation pattern also translated into changes in performance (Supplemental Table 1) but not in the expected manner. The highest F1-score was attained when the “ReLU” activation function was retained in the final layer (V_m_a_LeakyReLU_ReLU). To understand this paradoxical behavior, we examined the ground-truth labels of inactivated tiles (Figure 8). The “score-frequency” plot showed that most non-activated tiles were in the medium class. Thus, the default allocation of tiles to this class worked well for fold 2, which had the most medium damage tiles. After changing the activation function, the performance of other classes improved, as some of these tiles belonged to high or low damage classes.

**Figure 8.**
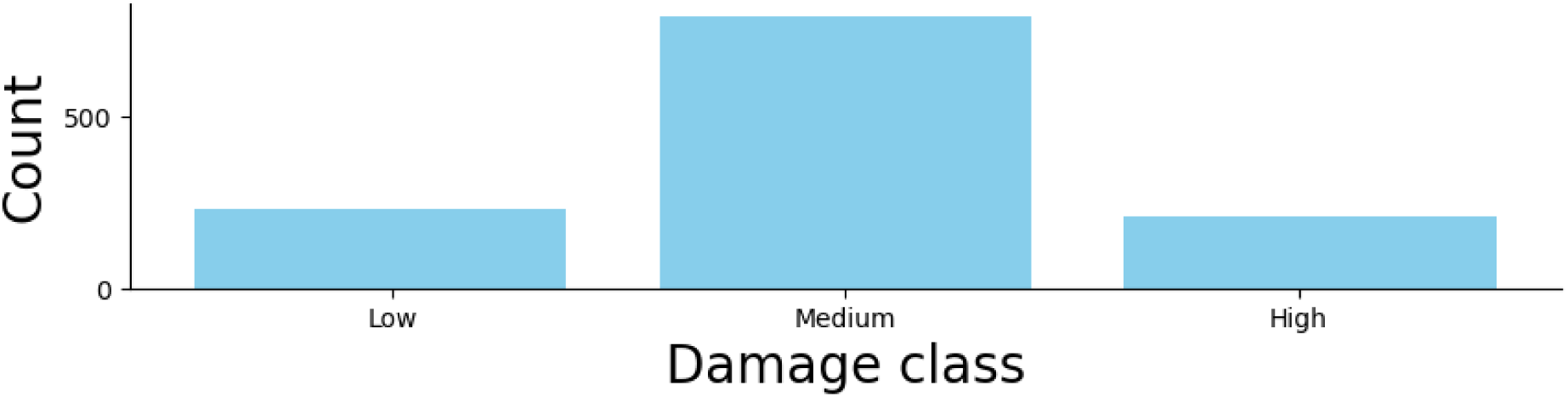
Count of non-activated tiles per class (V_r_a model Fold 2).

The primary motivation for employing Grad-CAM was to identify the features driving the model decisions. Upon further inspection of the activation heatmaps we observed that in mildly damaged tiles or tiles with fully visible airspace, the alveolar walls or the border between alveolar walls and airspace was highlighted. In contrast, in highly damaged tiles where airspaces were filled with inflammatory cells, the damaged areas (such as clumps of inflammatory cells) were highlighted. Due to Grad-CAM being a coarse map, establishing this connection was not entirely straightforward, but it seemed to hold true for almost all inspected tiles.

### SHAP

To show the importance of input features in the final model predictions, we calculated SHAP values [26] for the same subset of images as used in the Grad-CAM analysis (Figure 9). We first plotted SHAP value maps for all classes, where the classes are ordered from the most probable (left) to the least probable (right), as shown in Figure 9 Panel A. We then selected the most probable class predicted by the model and displayed the SHAP values for that class in Figure 9 Panel B.

**Figure 9.**
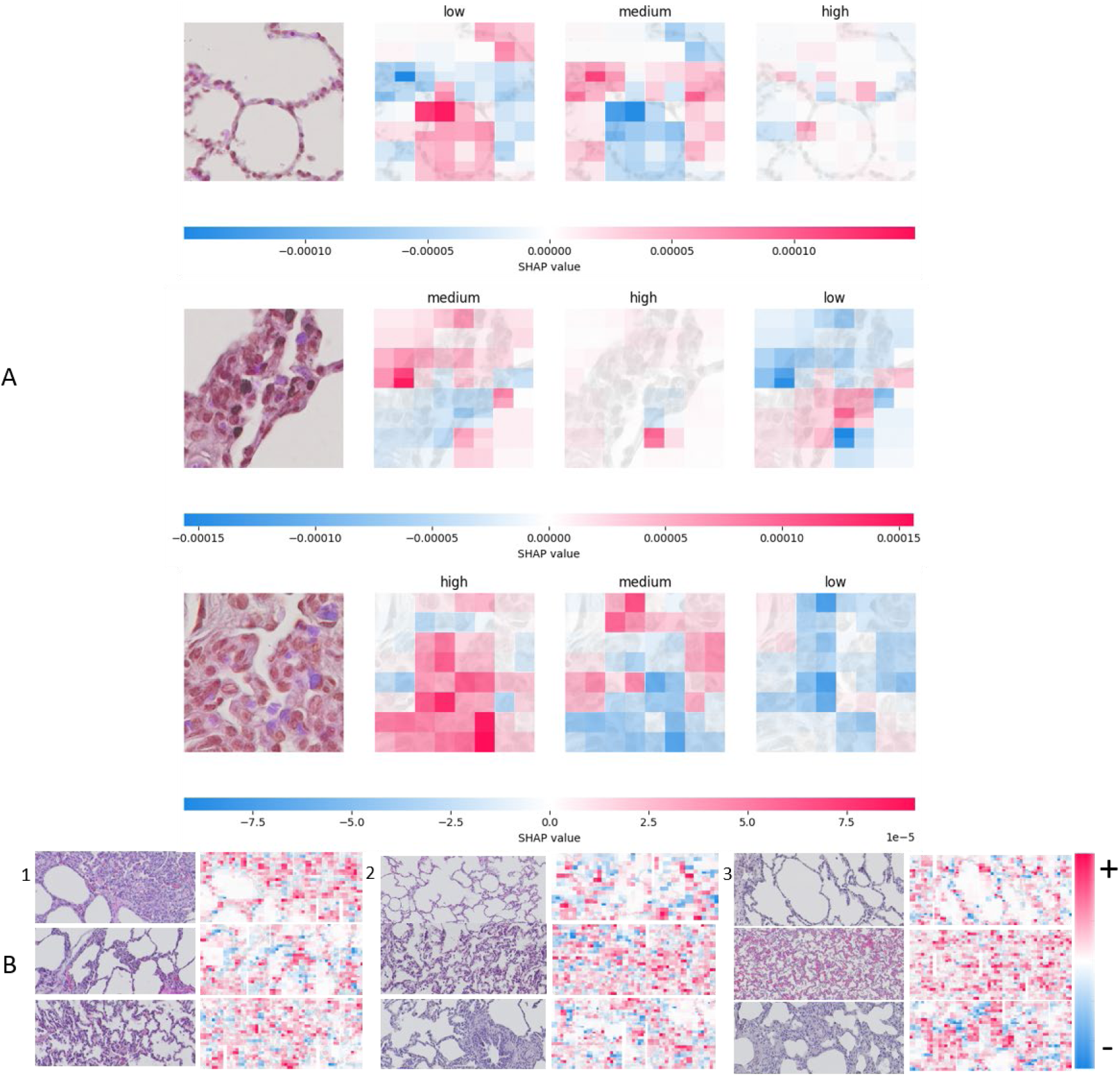
A) the SHAP values are arranged from the most probable predicted class (left) to the least probable (right) (e.g., the first row shows the image is mostly a low damage image). B) Shapley value plotted for sample images from “EN_m_a” model for tiles from folds 1,2 and 3. The most important features that have the highest contribution to the predicted class of each tile are depicted as red (most positive SHAP values) and blue (most negative values.

When examining image regions influencing a “high damage” decision, we saw that tissue areas showing pathological features such as clusters of inflammatory cells, fibroblasts occupying the lung airspace, or hemorrhage within the alveolar walls were predominantly associated with high positive SHAP values (red), indicating a strong positive contribution to the model’s decision for “high damage”. In contrast, healthy lung tissue was generally linked to the most negative SHAP values (blue), reflecting a negative contribution to predictions of “high damage”. Vice versa, in tiles classified as low or medium damage healthy areas were often labelled red. Additionally, in most tiles, background regions lacking tissue show SHAP values close to zero, indicating minimal influence on the model’s decision-making process.

### Deeper model fine-tuning: Effects on Model Performance and Generalization

As is common practice we used transfer learning [66] when training the initial models, meaning only the last layers were available for fine-tuning. We therefore investigated how training additional layers, which increases the number of trainable parameters, or the training of specific layer types, affected performance gains (Supplemental Table 2 and Figure 10, A and C). For the VGG16 base model, training an additional convolutional layer improved performance whereas adding two layers decreased both F1-score and accuracy. With limited data, convolutional may fail to capture robust features, causing activation maps to remain noisy or highlight irrelevant regions. Small datasets can also cause the model to memorize specific training samples rather than generalize, and some neurons may never activate properly, resulting in sparse activation maps. This effect is evident when inspecting the Grad-CAM heatmaps for the modified models (Figure 10B).

**Figure 10.**
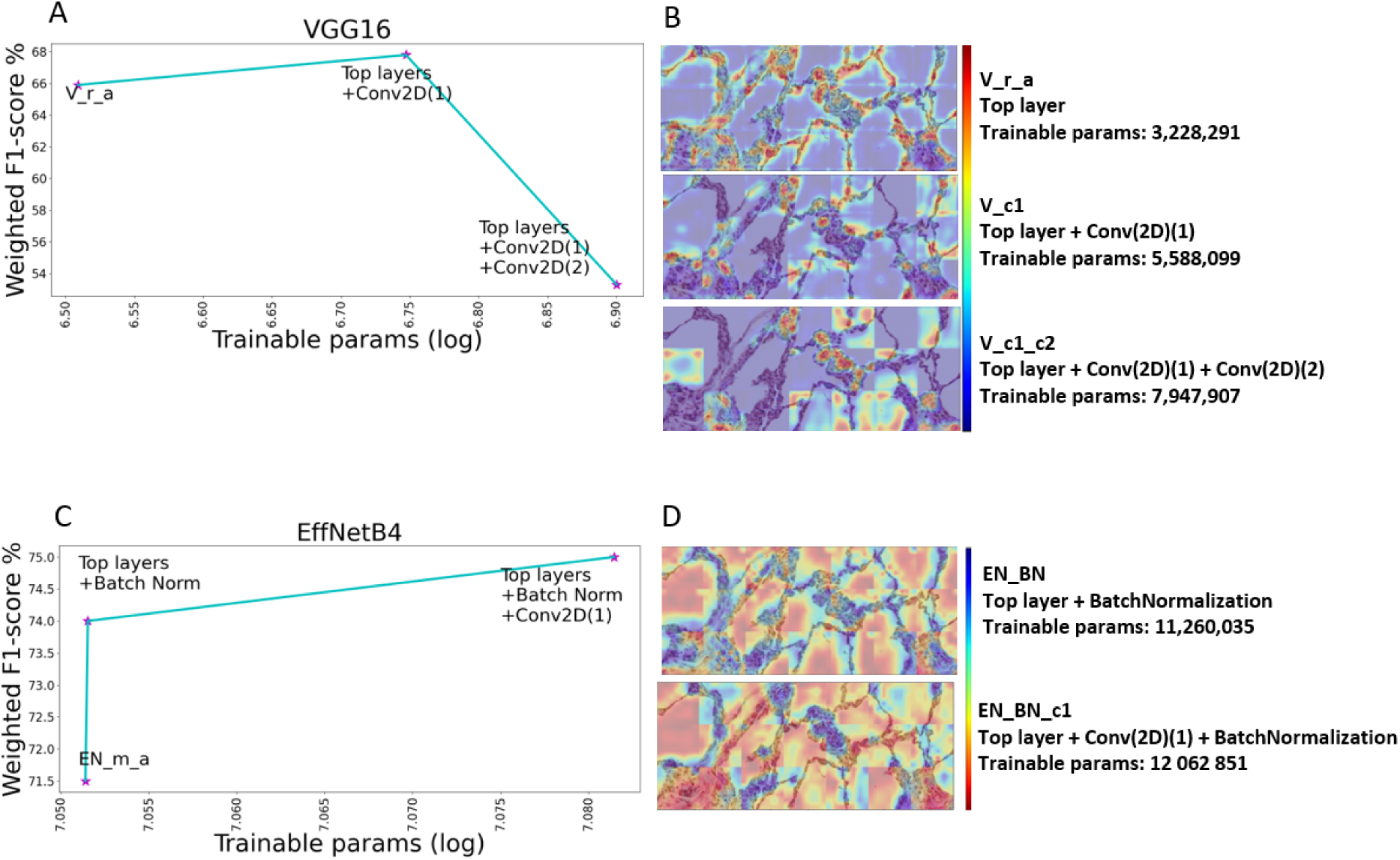
A, C) Relationship between model capacity and performance. The number of trainable parameters is plotted against the weighted-F1-score. B, D) Grad-CAM visualization of the last convolutional layer in the models that more layers are trained.

For the EffNetB4 model, training an additional batch normalization layer improved the F1-score with a further performance boost from also training a convolutional layer indicating a positive relationship between model parameters and performance at this scale. Retraining batch normalization (BN) layers stabilizes feature map distributions by normalizing activations across the batch, reducing extreme values and improving gradient flow. This leads to smoother, more stable activations, fewer spikes, and more effective learning in subsequent layers. Training BN often reduces the number of completely inactive neurons compared to un-normalized inputs. This effect is visible in the Grad-CAM heatmaps (Figure 10D).

### Analyzing Model Attention Using Convolutional Attention Layers

Next, we introduced an attention mechanism into the model architecture [6] (Figure 5) to determine the features the model focuses on during training and to directly assess the impact of specific features on predictions. The convolutional layers were utilized as filters to extract features from images that were then fed into the attention mechanism. Attention layers highlight important spatial or channel-wise features, making activation maps more focused and interpretable by emphasizing relevant regions while suppressing less important areas. Thereby, attention might also improve performance. Manual inspection of selected tiles revealed that the model mostly paid attention to the damaged parts of the images, especially inflammatory cells and collapsed airspaces, mainly along the tissue-airspace boundary. However, a slight reduction in overall performance across all classes was observed, particularly for the medium class. The sole exception was the high-damage class for which the EN_m_a_Attn model showed improvement.

**Figure 11.**
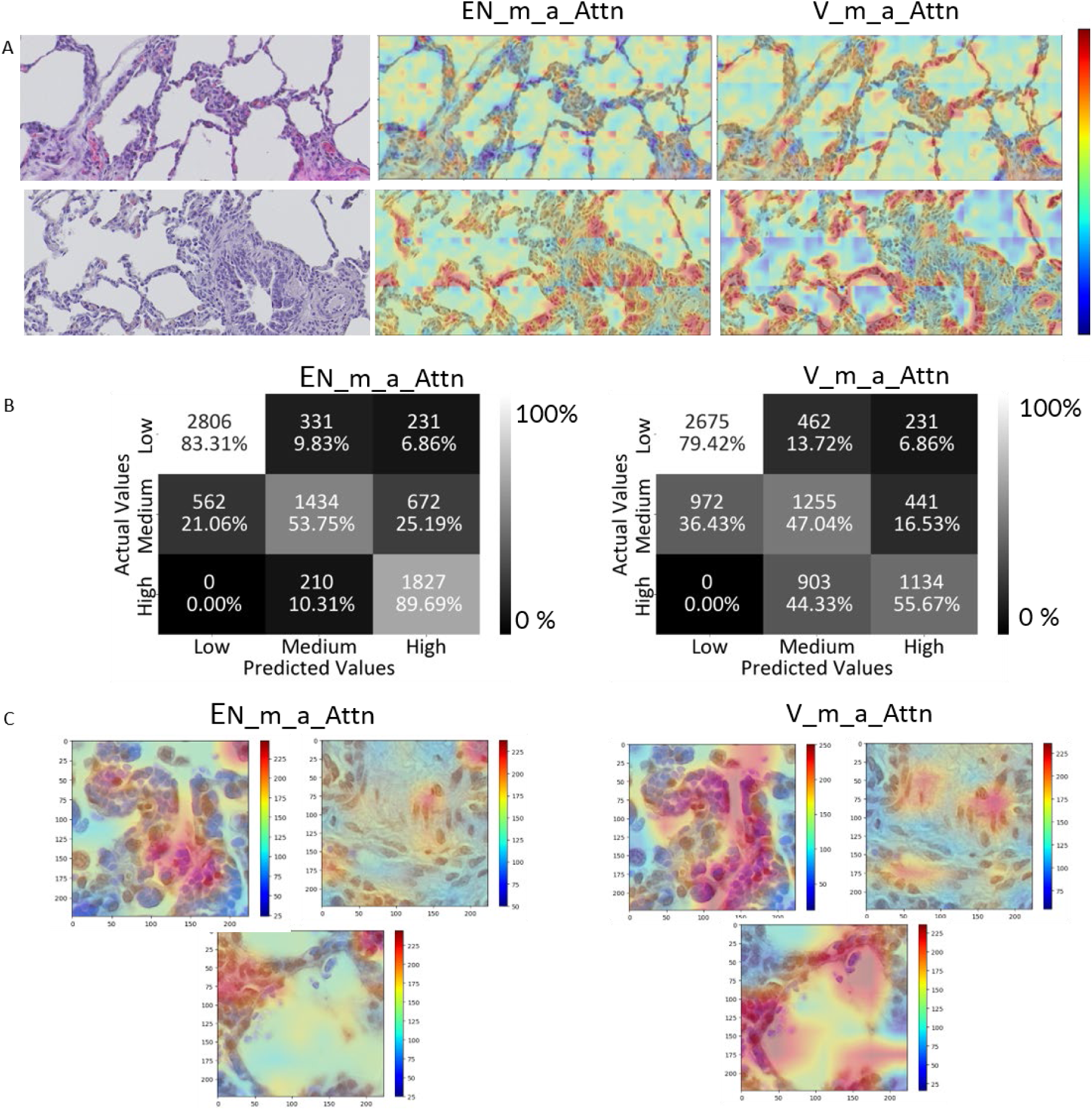
Impact of attention mechanism introduction. A) Attention maps indicating regions the model paid attention to. The red sections correspond to the highest attention weights. B) The confusion matrix illustrates the performance of the attention models across all three folds. C) Three example tiles where the attention model focused on damaged areas, specifically inflammatory cells and atelectasis or collapsed airspaces filled with fibroblasts.

### Statistical Analysis

To validate the statistical significance of the observed performance gains, we performed McNemar’s test. This relied on making predictions on all validation images with each of the improved models and comparing them to those from the respective baseline model (Figure 12A). V_r_a(Leaky_ReLU_ReLU) demonstrated the strongest improvement, achieving a 17.39% net accuracy gain over the baseline (p < 0.001), with improvement seen in over 27% of the samples. V_c1 and V_m_a_Attn also showed significantly improved accuracy of 10.05% and 7.18%, respectively (both p < 0.001). Conversely, V_c1_c2, which had a second layer unfrozen for training, showed significantly reduced performance (-15.66%, p < 0.001).

**Figure 12.**
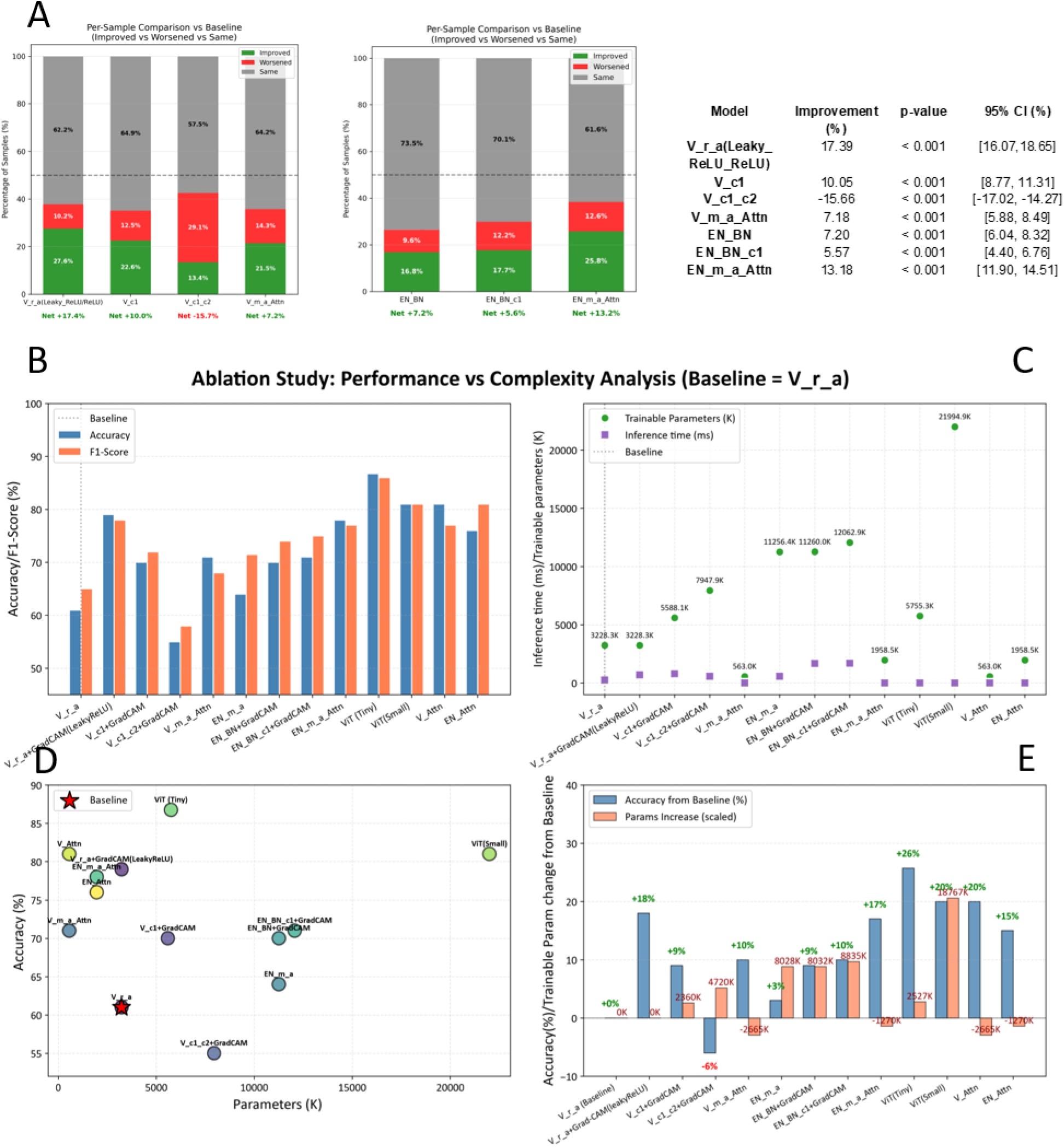
Performance Improvement vs Complexity. A) Pairwise differences between the modified models and their respective matching baseline (V_r_a or EN_m_a) relative to the true labels. Bars illustrate the percentage of cases in which the modified model performed the same, worse, or better compared with the baseline model. Statistical significance of the observed differences between the models were assessed by McNemar’s test with 95% bootstrap confidence interval (CI) B) Accuracy and F1-Score of all modified models. C). Comparison of trainable parameters and inference time across models with inference performed on a laptop GPU (see Methods). D) Visualization of the relationships between accuracy and trainable parameters. E) Bar chart showing the relative accuracy change (%) of each model alongside its relative change in trainable parameters compared to the V_r_a model.

All modified EN models demonstrated statistically significant improvements over the baseline (p < 0.001). EN_m_a_Attn achieved the strongest performance gain of 13.18%, EN_BN 7.20% and EN_BN_c1 5.57%. In summary, all observed changes were highly significant in McNemar’s tests, with bootstrap confidence intervals excluding zero.

#### Comparison with ViT Models

As we had seen a positive effect from the introduction of attention into the CNN architecture, we also evaluated ViTs, an alternative class of computer vision models that heavily rely on attention. Specifically, we chose to adapt ViT models that had recently been used successfully for binary classification of histopathology images from breast and gastric cancer datasets. The architecture was converted for 3-class classification, and two size variants, ViT (Tiny) and ViT (Small), were fine-tuned on our lung damage image dataset using the original training scripts. As the training procedure differed from that of our earlier CNN models, we trained a second set of our attention-modified CNN models alongside the ViTs using the same data processing and training procedure (V_Attn, EN_Attn).

Both the ViT models and the V_Attn and EN_Attn models performed with markedly higher accuracy and F1 score than the V_r_a and EN_m_a models from the earlier study (Table 3, Figure 12B, D, E), but a direct statistical comparison was impossible due to the different fold compositions. The ViT (Tiny) model reached the overall highest accuracy and F1 score, 87 and 86%, respectively, with the ViT (Small), V_Attn and EN_Attn models following closely behind (Table 3, Figure 12B, D, E).

**Table 3.** Comparison of ViT and matching CNN models.

| Device | Model | Parameters<br>Summary | Accuracy | F1-score | Inference<br>time/image |
| --- | --- | --- | --- | --- | --- |
| CUDA -<br>NVIDIA<br>GeForce<br>RTX<br>2060<br>CUDA<br>Version:<br>11.8<br>GPU<br>Memory:<br>6.44 GB | V_Attn | Total: 15,278,500<br>Trainable: **563,300**<br>Trainable ratio:<br>3.69% | 0.8059±0.0062 | 0.77±0.03 | 0.29±0.02ms |
|  | EN_Attn | Total: 19,508,908<br>Trainable: **1,958,500**<br>Trainable ratio:<br>10.04% | 0.7616±0.0100 | 0.81±0.05 | 2.55±0.15ms |
|  | ViT (Tiny) | Total: 5,755,331<br>Trainable: **5,755,331** | 0.8675±0.0100 | 0.86±0.03 | 2.25±0.15ms |
|  | ViT (Small) | Total: 21,994,883<br>Trainable:<br>**21,994,883** | 0.8133±0.0300 | 0.81±0.03 | 4.56±0.12ms |

When inspecting some of the ViT (Tiny) attention maps manually, we observed that areas with highest attention differed from those of the earlier V_m_a_Attn attention models (Supplemental Figure 2). However, we could not observe and immediately obvious alignment of the ViT (Tiny) attention maps with pathological features.

#### Multi-dimensional comparison of all trained models

To get a deeper understanding of the impact of the different model configurations and guide real-world decisions we compared all models across three dimensions: 1. Model performance as reflected in accuracy and F1-Score, 2. Model complexity as reflected in trainable parameter count, and 3. Computational cost as reflected in inference time (Figure 12B-E).

Rather than simply using XAI for explainability, a goal of our study was to convert the insights into model improvement. The original V_r_a model had a F1 score of 61% and an accuracy of 65%, while the EN_m_a model had a F1 score of 64% and an accuracy of 71.5% (Supplemental Table 2). With the exception of unfreezing a second convolutional layer (V_c1_c2), all our XAI-guided model modifications had improved performance (Figure 12A, B, D, E). The introduction of attention or the LeakyReLU activation function had the largest effects. Overall, the ViT (Tiny) performed best, with an 86% F1 score and 87% accuracy (Table 3, Figure 12B, D, E), albeit after having been trained and evaluated with a slightly different fold composition.

The number of trainable parameters varied greatly between our models, with an almost 40-fold difference between the ViT (Small) model and the models with least trainable parameters (V_Attn, V_m_a_Attn) (Figure 12B). In many cases, increasing model complexity leads to improved performance but we did not see any correlation between the number of trainable parameters and performance (Figure 12C, D). The smaller ViT (Tiny) model outperformed the ViT (Small) model, and several CNNs with much less trainable parameters were almost on par. Larger models, as well as some XAI approaches, are known to have increased inference time, which can hinder their in real-world applications. To assess the extend of this potential issue in our models, we measured inference times on a laptop with a RTX 2060 GPU (Table 3, 4). Of the attention-based models, the VGG16-based models (V_m_a_Attn and V_Attn) had by far the shortest inference time (around 0.3 ms/image), with an around 15-fold difference from the largest attention model ViT (Small) (around 4.6 ms). Still even the ViT inference times were dwarfed by the those seen with the other XAI approaches, SHAP and Grad-CAM, which ranged from around 100-1700 ms (Table 4, Figure 12C).

**Table 4.**
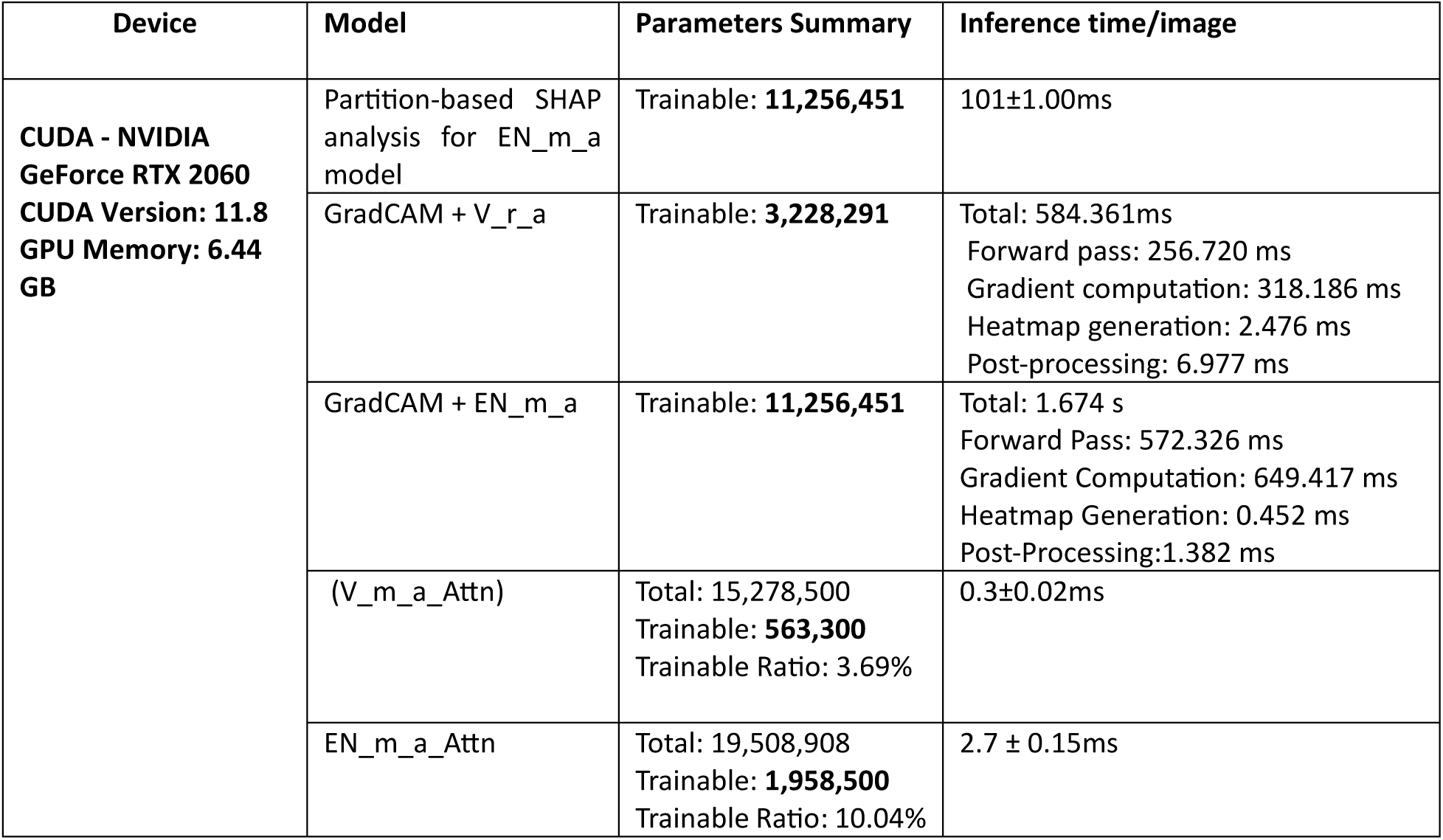
Comparison of model complexity and inference times among CNN models.

| Device | Model | Parameters Summary | Inference time/image |
| --- | --- | --- | --- |
| **CUDA - NVIDIA GeForce RTX 2060**<br>**CUDA Version: 11.8**<br>**GPU Memory: 6.44 GB** | Partition-based SHAP analysis for EN_m_a model | Trainable: **11,256,451** | 101±1.00ms |
|  | GradCAM + V_r_a | Trainable: **3,228,291** | Total: 584.361ms<br>Forward pass: 256.720 ms<br>Gradient computation: 318.186 ms<br>Heatmap generation: 2.476 ms<br>Post-processing: 6.977 ms |
|  | GradCAM + EN_m_a | Trainable: **11,256,451** | Total: 1.674 s<br>Forward Pass: 572.326 ms<br>Gradient Computation: 649.417 ms<br>Heatmap Generation: 0.452 ms<br>Post-Processing:1.382 ms |
|  | (V_m_a_Attn) | Total: 15,278,500<br>Trainable: **563,300**<br>Trainable Ratio: 3.69% | 0.3±0.02ms |
|  | EN_m_a_Attn | Total: 19,508,908<br>Trainable: **1,958,500**<br>Trainable Ratio: 10.04% | 2.7 ± 0.15ms |

## Discussion

XAI covers a variety of methods that have primarily been used to establish a connection between the features derived from the model and those reflecting medical phenotypes. Here, we integrated multiple XAI approaches into an iterative workflow, which not only focuses on model explainability but also reveals dataset and inference time limitations, and most importantly, directly translates insights into model improvements. As use case, we chose lung damage histopathology scoring, an understudied domain where XAI had not been applied to our knowledge. By improving the state-of-the-art model performance from our earlier study in this domain, we directly demonstrated the usefulness of our workflow.

Prior work has developed a series of XAI approaches and demonstrated their use in the histopathology domain [13]. In lung histology, post-hoc XAI methods have been used to understand features that drive model decisions, such as particular color frequencies or image regions [45–48, 50, 51] . Such features could then be related to medical characteristics. For example, Faria et al. [44] identified features by which the model differentiated between adenocarcinoma and squamous cell carcinoma using Grad-CAM and SHAP. Others have compared the features identified by different CAM methods [49]. Of particular interest among XAI methods are attention-guided architectures such as ViTs [51], which not only increase interpretability but in many cases also performance.

We built upon this work by integrating multiple XAI methods, including latent-space visualization [53], Grad-CAM [54], SHAP [26] and attention into a single workflow, but we widened the scope beyond pure model explainability. In addition to model explainability, the workflow also focuses on data explainability, using XAI approaches to reveal label inconsistencies and data distribution issues that affect model performance. Thereby our workflow can inform decisions about data acquisition and evaluation procedures. Most importantly, the workflow does not stop at explanation but instead translates the insights into XAI-informed model improvement. As last step, it investigates model complexity and inference time and relates them to performance, which can support decisions about real-world deployment.

Most prior work on XAI for lung imaging has focused on cancer. We therefore deliberately chose another less studied domain, lung damage scoring, in which XAI has not been applied to our knowledge. Specifically, we chose to apply our workflow to understand and improve models we recently trained for lung damage scoring of histology images from a porcine model of acute respiratory distress syndrome [52], selecting the two best individual models from this study (V_r_a and EN_m_a) for exploration.

Initially, we compared the model predictions and ground truth assessments provided by pathologists using unsupervised clustering of latent representations. While difficult to make a final assessment it nevertheless appears that model predictions may separate classes more clearly and align better with treatment groups and tissue damage visible at macroscopic level. Our earlier study [52] already hinted at how difficult it is even for well-trained images to score these images, with relatively poor inter-annotator and intra-annotator agreement. It is thus possible that the model predictions were more correct than the ground truth labels. Our investigation highlighted a challenge in medium-class tiles, where human classification often leans towards high or low damage.

Subsequently, we employed passive, post-hoc XAI algorithms such as Grad-CAM and SHAP values. Prior studies have shown their usefulness in histopathology by helping to reveal image areas and features that inform model decision-making [43–51]. Many of these studies where performed on cancer and matching control samples but to our knowledge none on lung damage scoring. Here we demonstrated that the XAI approaches are equally useful in this domain, with the models identifying damaged regions. This was particularly evident in highly damaged tiles, where the model highlighted the affected regions and pathological features. In addition, features associated with lower levels of damage—characterized by the absence of clear injury indicators such as larger air spaces and thinner alveolar walls—were frequently the most strongly emphasized in Grad-CAM heatmaps and high-magnitude SHAP values. This suggests that tiles characterized by larger airspaces, typically reflecting thinner alveolar septa, fewer inflammatory cells, or an absence of additional injury markers such as hyaline membrane formation were correctly identified as “low damage”.

Guided by these insights, we explored the unfreezing of additional layers, changes in activation function and the introduction of attention mechanisms. Increasing model complexity (i.e. adding trainable parameters) is generally beneficial, provided there is enough data. However, in our case, as in many histology-related settings, the dataset was very small. This could be the reason that ViT (Tiny) outperformed the larger ViT (Small), and that unfreezing a second convolutional layer in the VGG16 architecture substantially reduced performance. However, the type of layer that is unfrozen appeared to also matter as unfreezing both a convolutional and a batch normalization layer in the EfficientNet architecture increased performance, rather than reducing it.

Attention can be a means to improve performance as well as explainability. We also observed this. All CNN architectures with attention showed improved performance compared to the respective base models (V_r_a and EN_m_a). In addition, the best performing model was the attention-based ViT model. One caveat with the ViT models is that the fold composition of these models differed, because the original training script was retained to ensure better comparability with prior work. With a dataset this small, it may have influenced the difficulty of the evaluation and thus slightly improved the performance scores, especially since the fold composition algorithm randomly distributes tiles rather than keeping all related images from the same pig together like our fold composition algorithm did. This notion is supported by the observation that the second set of attention-modified CNN models (V_Attn, EN_Attn), which was trained in the same manner as the ViT models, also scored slightly higher than the first set (V_m_a_Attn, EN_m_a_Attn), which was trained on the original fold composition. Still, it seems clear overall that introducing attention in some form is beneficial for this task.

When it comes to real-world use, performance is often not the only concern. Inference time governs computational cost and image throughput. The ViT models had the highest model complexity, as measured in trainable parameters, which translated into a markedly increased inference time compared to the attention modified VGG16 models. As the performance of the V_Attn model was only slightly lower, the smaller faster models may thus be preferable in many real-world settings. Perhaps the results would have favoured the ViT models more strongly, had the dataset been larger, but unfortunately small datasets are an all too common limitation in the medical domain.

A much larger issue with inference time was demonstrated for the other XAI approaches, GradCAM and SHAP, with the former elevating inference times to up to 1.7 s per image. In many cases, such long inference times, may indeed prevent these techniques from being feasible during eventual deployment. However, they could still be used during model development, where we confirmed their usefulness even within the domain of lung damage scoring. GradCAM in particular had revealed that many tiles were not activated for the predicted class with the original model and thus did not contribute to the prediction. This was likely related to the "dying ReLU" problem, where neurons become inactive. Replacing the ReLU activation function indeed improved the activation patterns. However, the situation was a bit more complex with regards to performance. The best model (V_r_a_ReLU_LeakyReLU) of this experimental series had indeed a different activation function configuration than the original, but it still had many non-activating tiles as it retained the ReLU function in the last layer. Models with other activation functions showed much better activation patterns but did not have higher performance scores overall. This was perhaps because overall performance was somewhat skewed by the fact that the default predicted class was also the most common class, and that this class was especially common in one of the folds. Entirely equal fold distribution was not possible because the algorithm also considered the relatedness of images. Future work, ideally with larger image datasets, should dive deeper into this question. The present study clearly demonstrated how its XAI workflow can reveal important issues with activation functions and datasets that can guide immediate model improvement as well as inform decisions about data acquisition and evaluation design.

In summary, the contribution of this study is not the introduction of a new explainability algorithm, but the integration of complementary XAI approaches into an iterative workflow that can guide model development and deployment within digital pathology. Latent space analysis is used to identify potential label inconsistencies, Grad CAM and SHAP are used to assess whether predictions are based on pathology relevant regions, and these findings are then used to guide targeted model refinement through fine tuning and attention mechanisms. Assessments of model parameters and inference times are related to performance to guide further development and real-world deployment decisions. As a consequence of applying our workflow we also improve the state of the art of lung damage scoring.

## Conclusion

While many deep neural networks with CNN or ViT architectures perform well in histology image analysis, bridging the gap between pathology and model outputs remains non-trivial. Different interpretability algorithms have inherent limitations and operate at distinct representational levels capturing different aspects of model behavior. While some methods primarily emphasize spatial localization of salient regions, others focus on feature attribution or the separation of class-specific representations. Consequently, these approaches may provide complementary rather than identical insights into the underlying decision-making processes of the model.

Nevertheless, XAI interpretation is not straightforward and human expertise in the loop is still essential. To achieve more reliable and pathologist-convincing results, it may also be necessary to define more granular, pathology-specific features rather than relying on broad combined categories (e.g., low, medium, high damage). As XAI methods also add computational expense and analytical complexity their benefits must also be weighed against explainability needs in every scenario.

Our workflow addresses these demands. It combines multiple XAI approaches in a meaningful way to gain better understanding of data and models, and translates them into better performing models. It also highlights areas of uncertainty and real-world trade-offs between performance, model complexity and inference times. This makes it a practical template that can help improve transparency as well as guide model development, data acquisition, evaluation design and deployment.

#### Box 1. Strategies to increase transparency and explainability of models for histology analysis

Reporting of EDA findings regarding dataset sample composition, biases and flaws

- Image characteristics discovered by manual inspection
- Label distribution and correlation between annotators
- Dataset size and distribution of number of images per sample group

Publication of visual examples of augmentations

Reporting of training/evaluation/test dataset selection procedure and composition

Reporting of data pre/postprocessing and model architecture

Reporting of training strategy (e.g. training schedule, covered hyperparameter space) and evaluation metrics

Discovering Structure in Latent Representations via Unsupervised Clustering

Gradient-weighted Class Activation Mapping (Grad-CAM)

SHapley Additive exPlanations (SHAP)

Deeper Training and evaluating effects on model performance

Attention introduction and analysis of attention maps

## Ethics approval and consent to participate

Not applicable

## Consent for publication

Not applicable

## Authors’ contributions

Conceptualization/Methodology: SKR (primary), SA

Data curation/Formal analysis/Investigation/Software/Validation: SKR

Funding acquisition/Project administration/Supervision: SA

Writing – original draft: SKR, SA

Writing – review & editing: SKR, SA, MN

## Acknowledgements

We thank all members of the “Lung Bioengineering and Regeneration” group from Lund University for helpful comments throughout the development of this project and the dataset. We thank Jacob Vogel and Lijun An from “Neurodegenerative Research” group, Lund University, for critical review and helpful comments on the manuscript.

We also acknowledge the following research environments and networks which support our work: AI Lund, AIR Lund, Lund University Profile Area “Nature-based Future Solutions”, Lund University Profile Area “Natural and Artificial Cognition”, Lund University Profile Area “Proactive Ageing”, LTH Profile Area “AI and Digitalization”, LTH Profile Area “Engineering Health”, Strategic Research Area “EpiHealth”, Strategic Research Area “eSSENCE”, and PhenoTarget.

## Supplemental Materials

**Supplemental Figure 1.**
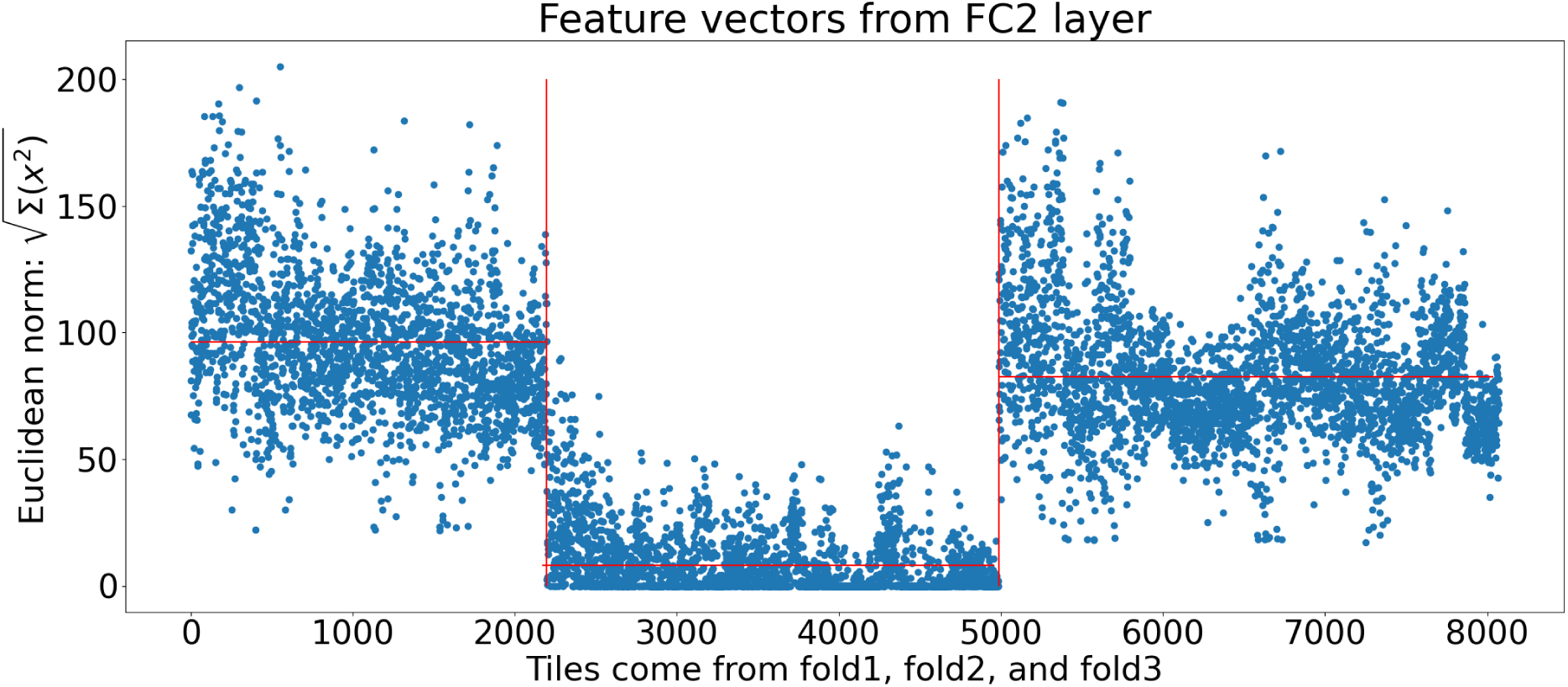
Euclidean norm of latent space vectors (all folds) for each tile extracted from the fully connected layer of the V_r_a model.

**Supplemental Table 1.** Comparison of performance of “V_r_a” models. The table shows precision, recall, and F1-score for all folds for different activation functions for the last two fully connected layers. The rows show the “V_r_a” models with “ReLU”, combination of “LeakyReLU-ReLU”, “ELU” and “LeakyReLU” activation functions.

|  |  | Fold1 |  |  | Fold2 |  |  | Fold3 |  |  | Average |  |  |
| --- | --- | --- | --- | --- | --- | --- | --- | --- | --- | --- | --- | --- | --- |
|  |  | Precision | Recall | F1-score | Precision | Recall | F1-score | Precision | Recall | F1-score | Precision | Recall | F1-score |
| V_r_a (ReLU) (B) | Micro average | 0.85 | 0.85 | 0.85 | 0.8 | 0.80 | 0.8 | 0.5 | 0.5 | 0.5 | 0.72 | 0.72 | 0.72 |
|  | Weighted average | 0.88 | 0.85 | 0.87 | 0.81 | 0.8 | 0.80 | 0.89 | 0.5 | 0.64 | 0.86 | 0.72 | 0.78 |
| V_r_a (LeakyReLU) (B) | Micro average | 0.75 | 0.75 | 0.75 | 0.72 | 0.72 | 0.72 | 0.86 | 0.86 | 0.86 | 0.78 | 0.78 | **0.78** |
|  | Weighted average | 0.83 | 0.75 | 0.79 | 0.82 | 0.72 | 0.76 | 0.87 | 0.86 | 0.86 | 0.84 | 0.78 | **0.81** |
| V_r_a<br>( ' ELU' ) | Micro average | 0.75 | 0.75 | 0.75 | 0.72 | 0.72 | 0.72 | 0.63 | 0.63 | 0.63 | 0.70 | 0.70 | 0.70 |
|  | Weighted average | 0.83 | 0.75 | 0.79 | 0.82 | 0.72 | 0.76 | 0.82 | 0.63 | 0.71 | 0.82 | 0.70 | 0.76 |
| V_r_a<br>( ' Leaky ReLU' ) | Micro average | 0.74 | 0.74 | 0.74 | 0.55 | 0.55 | 0.55 | 0.85 | 0.85 | 0.85 | 0.72 | 0.72 | 0.72 |
|  | Weighted average | 0.82 | 0.74 | 0.78 | 0.76 | 0.55 | 0.64 | 0.93 | 0.85 | 0.89 | 0.84 | 0.72 | 0.77 |

**Supplemental Table 2.** Characteristics of the investigated models. The table shows the base model from which additional layers were trained, the types of trained layers, the number of parameters added to the original model (where only the top layers were trained) total trainable parameters, the accuracy of the models, and the F1-score of the models, averaged across all folds. Rows 1 and 5 have already been included in [52]

|  | Base Model | Trained layers | #Added parameters | #Trainable parameters | Accuracy | F1-score |
| --- | --- | --- | --- | --- | --- | --- |
| 1 | **EfficientNetB4 (EN_m_a) [52]** | Top layers (Figure 3_A) | 0 | 11,256,451 | 64% | 71.5% |
| 2 | **EfficientNetB4 (EN_BN)** | Top Layers + Batch Normalization | $\frac{7\ 168}{2} = 3584$ | 11,260,035 | 69% | 74% |
| 3 | **EfficientNetB4 (EN_BN_c1)** | Top Layers + Convolution (1) + Batch Normalization | $\frac{7\ 168}{2} + 802,816 = 804,400$ | 12,062,851 | 69% | 75% |
| 4 | **VGG16 (V_r_a) [52]** | Top Layers (Figure 3_A) | 0 | 3,228,291 | 61% | 65% |
| 5 | **VGG16 (V_c1)** | Top Layers + Convolution (1) | 2,359,808 | 5,588,099 | 71% | 72% |
| 6 | VGG16<br>(V_c1_c2) | Top Layers +<br>Convolution (1) +<br>Convolution<br>(2) | $2 \times 2,359,808$<br>$= 4,719,616$ | 7,947,907 | 55% | 58% |

**Supplemental Figure 2.**
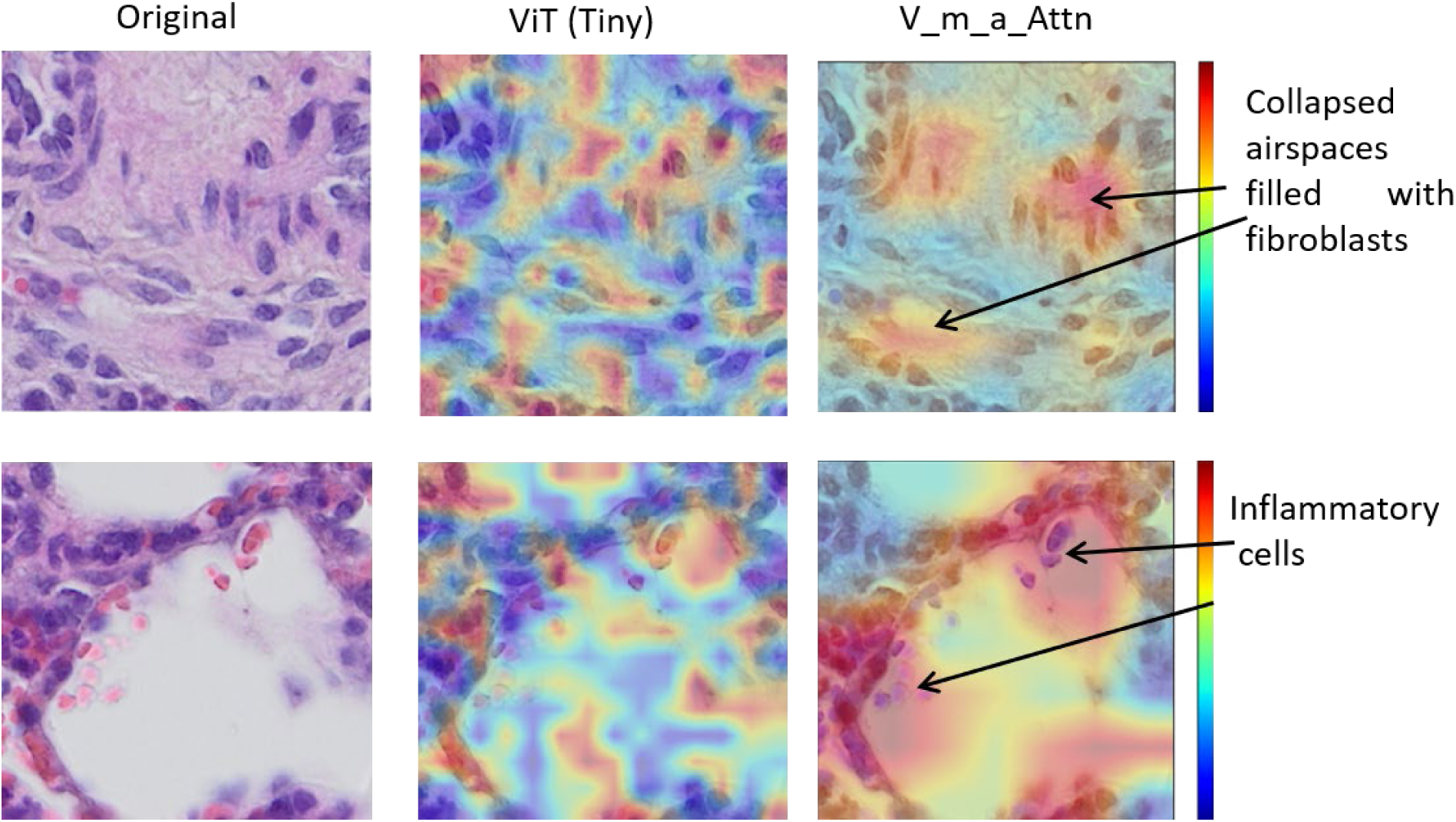
Comparison of attention maps between the ViT (Tiny) and V_m_a_Attn model.

## Declaration of generative AI and AI-assisted technologies in the manuscript preparation process

During the preparation of this work, the authors used ChatGPT and DeepSeek to improve grammar, revise selected sentences, and enhance the clarity of algorithm descriptions. ChatGPT, Claude and Gemini Flash (Extended Thinking) were also used to critique the manuscript structure. ChatGPT was also used to assist in conceptualizing the graphical abstract. The graphical abstract itself was created entirely by the authors **without** the use of generative AI. The authors reviewed and edited the output as needed and take full responsibility for the content of the published article.

## Abbreviations

ARDS: Acute Respiratory Distress Syndrome
CNN: Convolutional Neural Networks
ECMO: Extracorporeal Membrane Oxygenation
Grad-CAM: Gradient-based Class Activation Mapping
LPS: Lipopolysaccharides
MV: Mechanical Ventilation
PCA: Principal Component Analysis
SHAP: Shapley Additive exPlanations
t-SNE: T-Distributed Stochastic Neighbor Embedding
UMAP: Uniform Manifold Approximation and Projection
ViT: Vision Transformer
XAI: Explainable AI

